# Movement mechanisms explain current animal density patterns and predict their reorganization under global change

**DOI:** 10.64898/2026.10.05.756739

**Authors:** Robin Bauknecht, Will Rogers, Aaron T. Pearse, Walter Jetz

## Abstract

**Aim:** To test whether individual movement mechanisms inferred from animal tracking data can be scaled up to explain landscape-scale patterns of population density and improve predictions of population redistribution under future environmental change.

**Location:** Texas Gulf Coast, USA

**Time Period:** 2009–2018

**Major Taxa Studied:** Whooping Crane (*Grus americana*)

**Methods:** We parameterized individual-level step-selection functions using 14,898 GPS locations from 22 Whooping Cranes and propagated the resulting movement and habitat-selection mechanisms across the landscape to derive steady-state distributions of space use. We compared these movement-informed predictions with correlative species distribution models fitted to the same data. Predictions were evaluated using withheld tracking observations and fully independent population-level aerial-survey data. We then projected both approaches under sea-level-rise scenarios to assess how future habitat loss and fragmentation alter crane space use.

**Results:** Movement mechanisms derived from individual-level tracking data captured the mechanistic dependence of population density on habitat quality and spatial configuration. When projected across the landscape, these mechanisms reproduced fully independently observed density patterns and outperformed conventional species distribution models. Because predictions arose from movement mechanisms rather than occurrence-environment correlations, movement-informed models also captured the effects of habitat reconfiguration under sea-level rise. They projected pronounced population redistribution and an expansion of core space-use areas of up to 80% which conventional models missed.

**Main Conclusions:** Individual movement mechanisms derived from biologging data can scale up to explain current population structure and predict future redistribution under environmental change. By mechanistically accounting for how habitat configuration constrains accessibility, movement-informed models reveal severe impacts of fragmentation that are missed by conventional distribution models. Rapidly expanding biologging datasets therefore offer a scalable basis for connecting individual behavior to macroecological patterns and improving biodiversity forecasts in changing landscapes.

## Introduction

Human activities are reshaping landscapes worldwide, causing the loss and fragmentation of natural areas (Haddad *et al*. 2015; Hansen *et al*. 2013; Venter *et al*. 2016). These changes to the size and arrangement of habitats fundamentally alter how animals can use space, requiring them to navigate complex mosaics to persist (Brodie *et al*. 2025; Fletcher Jr. *et al*. 2019). Understanding and predicting animal space use in increasingly fragmented ecosystems is thus central to ecology and conservation (Guisan *et al*. 2013; Pressey *et al*. 2007). The importance of patch size, connectivity, and quality for species persistence has long been emphasized in metapopulation theory (Hanski 1999; Thomas *et al*. 2001), which formalized how extinctions and colonization dynamics generate occupancy patterns across fragmented ecosystems. While these frameworks have been highly successful in shaping our understanding of patch occupancy and persistence, they treat behavior as a population-level rate governing dynamics between discrete patches (Hanski 1999, 1998). As a result, these models leave unresolved how fine-scale movement decisions of individuals navigating heterogeneous landscapes give rise to the continuous spatial variation in population density observed within and across patches.

Animal space use is inherently dynamic and shaped by how individuals move through, perceive of, and respond to complex landscapes (Mueller & Fagan 2008; Nathan *et al*. 2008). To persist in fragmented or temporally varying environments, animals continually make movement decisions, balancing access to high-quality habitat with the costs and constraints of navigating space (Fig. 1A). These small, repeated choices accumulate through time, potentially concentrating use in larger, better-connected, or more centrally located areas while leaving others underused despite their apparent suitability (Fig. 1B). Thus, space use is not a simple reflection of where suitable habitat occurs but rather the outcome of cumulative animal behavioral responses to spatial structure given movement constraints (Moorcroft *et al*. 2006; Potts & Börger 2023). Across many individuals, these recurrent patterns may aggregate to yield spatial variation in what is recognized as population density (i.e., the number of individuals in a given area; Fig. 1C). If so, population distributions and densities may arise as emergent properties of individual movement and habitat-selection decisions. Consequently, population redistribution under environmental change should also emerge from how these mechanisms interact with shifting landscape structure (Fig. 1D). Individual-level movement and selection mechanisms thus could provide a generative basis for predicting how populations organize – and reorganize – across landscapes.

**Figure 1:**
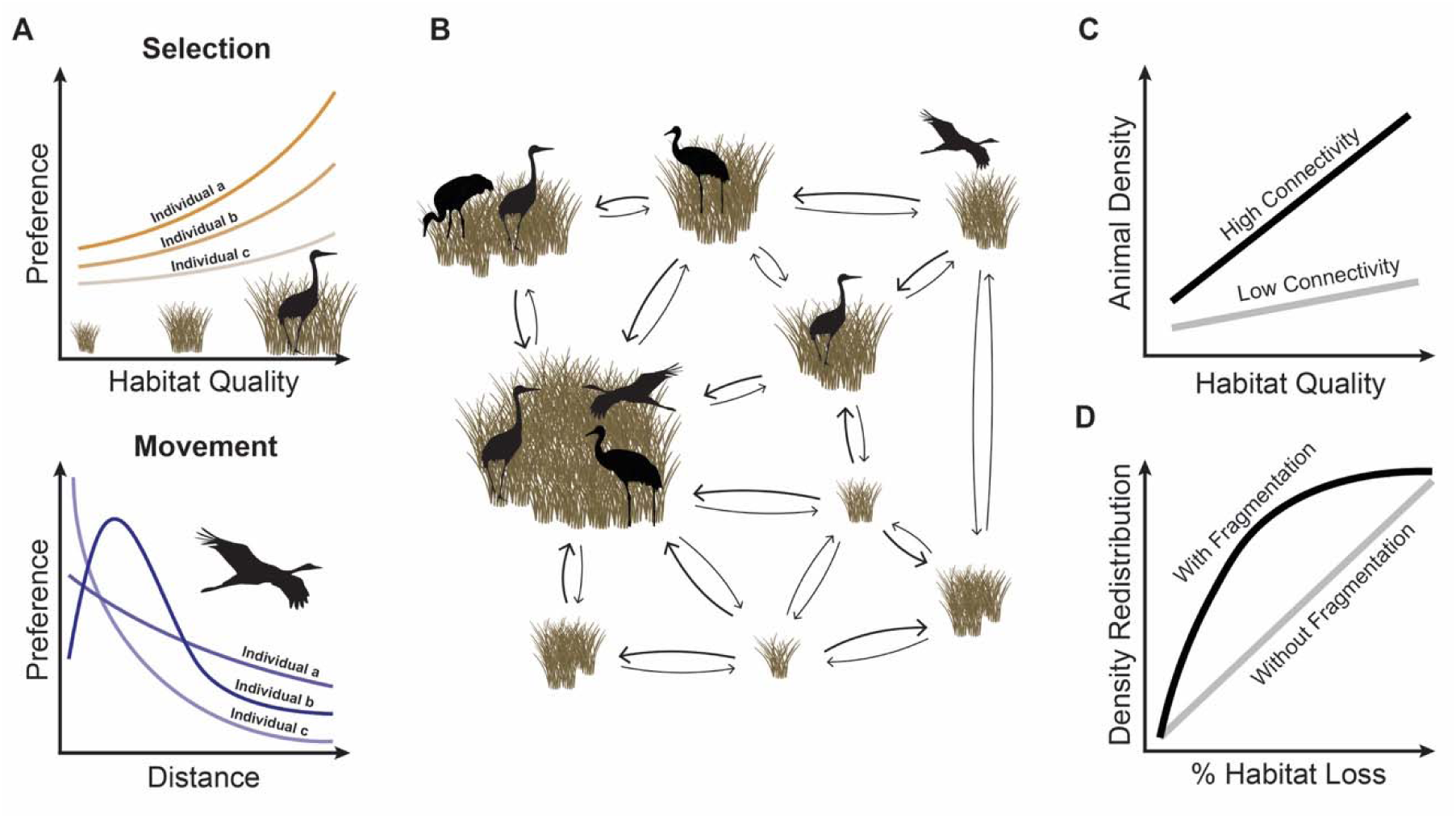
Individual movement and selection scale up to population-level spatial patterns. (A) Animals balance the benefits of using high-quality habitat against the costs of moving through space, generating preferences for both habitat quality and movement distance. (B) These local movement mechanisms shape individual-level space use across heterogeneous and spatially structured landscapes and, when aggregated across individuals, give rise to emergent population-level spatial structure. (C) This produces density patterns in which high-quality habitat accumulates more individuals when it is well connected than when it is spatially isolated. (D) As habitat is lost, changes in habitat configuration – such as fragmentation – therefore amplify spatial redistributions of populations relative to predictions that ignore spatial structure.

At present, these considerations of spatial structure and movement constraints stand in contrast to how ecological and conservation research typically conceptualize species distributions. Species distribution models (SDMs) in a broad sense treat distributions as static projections of a species’ realized environmental niche onto the landscape, often relying on the simplifying assumption that suitable habitat is broadly accessible and occupied according to suitability (Elith & Leathwick 2009; Guisan & Zimmermann 2000; Soberón 2007). These correlative approaches can describe where habitat occurs, but by design SDMs alone cannot capture the behavioral mechanisms that govern the accessibility and ultimate use of habitats. As a result, even with dedicated covariates for habitat structure (e.g., FRAGSTATS; McGarigal 1995), such models at best partially capture the cumulative effects of connectivity constraints and accessibility gradients that structure real populations, and are therefore not expected to capture resulting patterns of space use and emergent population density (Avgar *et al*. 2016; Rogers *et al*. 2025). Recently, animal movement data has emerged as an avenue for addressing the interaction of animal behavior and spatial structure through tools such as ergodic movement frameworks (Ellison-Neary *et al*. 2026; Rogers *et al*. 2025) and agent-based simulations (Michelot *et al*. 2019; Potts & Börger 2023). These approaches empirically derive local movement and habitat selection rules, propagating behavioral decisions through landscape structure to recover the stabilized patterns of space use. By framing distributions as the ergodic consequence of movement behavior, they provide a direct, testable way to evaluate whether observed density gradients emerge from individual movement mechanisms – and whether modeling these mechanisms yields more accurate predictive maps of population density.

The conceptual distinction of space use as emerging from the interplay of habitat structure and behavioral constraints has potentially profound implications for understanding biodiversity responses to anthropogenic change. This is because changes in the environment act on biodiversity through two broad mechanisms: alterations in the amount of habitat available (the area reflecting a species’ niche) and changes to its spatial configuration (Fahrig 2017; Hanski 2011). However, neither metapopulation dynamics nor correlative distribution modeling can truly address changes in the spatial configuration of habitats. Changes in habitat configuration reshape the pathways, bottlenecks, and bridging patches that govern how animals move within and between habitats (Brodie *et al*. 2025; Urban & Keitt 2001). As connectivity erodes or is rerouted, centers of attraction can shift abruptly, with suitable but newly isolated areas losing influence and previously peripheral locations possibly gaining prominence (Urban & Keitt 2001). Those changes should thus strongly affect the cumulative movements of organisms in the landscape and, in turn, the resulting population density distribution (Fig. 1D). Theory suggests that such behavioral redistribution effects could be nonlinear with modest changes in habitat configuration producing outsized differences in population density (Bascompte & Solé 1996; Swift & Hannon 2010). Without accounting for habitat accessibility and its effects on animal movement, modelling approaches are unable to anticipate how populations will spatially redistribute, split, or collapse as landscapes change.

Here, we test whether population-level patterns of animal space use and density can be understood and predicted as emergent outcomes of individual movement behavior responding to landscape structure. Using the wintering range of the endangered *Grus americana* as a case study, we evaluate whether such movement mechanism-informed predictions outperform correlative approaches in explaining population density and in forecasting how populations redistribute as habitat is lost and spatially reconfigured. More broadly, we demonstrate how empirically derived movement mechanisms provide a predictive foundation for linking individual behavior to present-day population structure and for projecting population reorganization in complex and changing landscapes.

## Methods

### Steady-state distributions (SSD)

We applied the SSD framework (Rogers *et al*. 2025) to model Whooping Crane space use under the combined influence of habitat selection and movement (Fig. S1). SSD uses eigenvector decomposition of transition matrices parametrized using step selection functions (SSF) to derive steady-state distributions of animal space use across landscapes (Fletcher Jr. *et al*. 2019; Rogers *et al*. 2025). Landscapes are represented as surfaces of discrete occupancy states between which transitions occur with defined probabilities. For a landscape G of n cells, transition probabilities between all n cells can be summarized in a stochastic matrix A of dimension n x n. Element A_i,j_ in this matrix contains the probability for an animal to move from cell G_i_ to cell G_j_. The probabilities of moving from a cell G_i_ to any other cell G_j_, including staying in G_i_, must sum to one. Thus the matrix is row-stochastic:

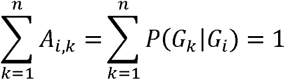

Matrix A can be empirically parameterized through movement-based integrated step selection functions (iSSFs; Avgar *et al*. 2016), specifically by computing the relative strength of selection (RSS) for each possible movement (Avgar *et al*. 2017). RSS quantifies selection by comparing a given movement to a reference condition and is defined for movement from G_i_ to G_j_ as:

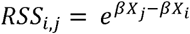

where X_j_ and X_i_ are vectors of environmental and movement covariates describing target cells G_j_ and reference cell G_i_, respectively, and β is a vector of selection coefficients estimated from the SSF. RSS is converted to movement probabilities by normalization over destination cells:

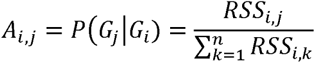

This yields movement probabilities conditional on starting in G_i_. Restricting movement to a defined maximum distance around each focal cell (here, 99th percentile of observed step lengths; Rogers *et al*. 2025) makes A sparse. This reduces computational cost while preserving realistic movements. Finally, we find the first principal eigenvector of the row-stochastic matrix A, describing the steady-state distribution of animal space use in G, hence “SSD” (see Rogers *et al*. 2025 for details).

### Whooping Crane space use modelling

#### Tracking data

Tracking data was sourced from an existing dataset containing tracking information for 68 individuals tagged between 2009 and 2018 (Pearse *et al*. 2020). The data was filtered to retain locations within the study area (≈ 90 x 90 km in coastal Texas, USA, Fig. S2) and remove records with missing coordinates or timestamps. Locations with velocities exceeding 2.6 m/s were classified as in-flight (following Lehnen *et al*. 2024) and excluded.

#### Environmental predictors

Environmental predictors were derived from the NOAA Marsh Migration dataset from 2016 (https://coast.noaa.gov/slrdata/), which provides land cover distributions for both current conditions and projected future sea level rise scenarios. The dataset’s land cover classification without sea level rise is directly sourced from the NOAA Coastal Change Analysis Program (C-CAP; National Oceanic and Atmospheric Administration, Office for Coastal Management 2016), a dataset previously proven to be effective in Whooping Crane habitat suitability modelling (Lehnen *et al*. 2024; Metzger *et al*. 2020). The dataset classifies land cover into 13 categories at a 10 m resolution. To ensure that individual movement tracks include locations across all categories, the classes were aggregated into four broader categories: upland, palustrine wetlands, estuarine wetlands, and open water (Fig. S2). Additionally, a layer of distance to shoreline (approximated by distance to the next pixel of type “Unconsolidated Shore”) was derived from the original classification. To reduce computational cost, land cover layers were resampled to 150 m resolution by calculating the proportion of each 10 m land-cover class within each 150 m grid cell. This resolution represented a compromise between retaining landscape heterogeneity and computational feasibility, with all modelling approaches using the same environmental layers for comparison. To avoid scale-driven biases during model fitting, all environmental predictor layers were rescaled to a common range.

#### Model development

For each individual, GPS locations were split chronologically into training (75%) and evaluation (25%) sets (with similar distributions of movement characteristics across sets, see Fig. S12). The locations were converted into movement tracks and resampled to the median GPS transmission interval, which was either 5 or 6 hours, depending on the individual. Only individuals with > 300 training steps were then retained to ensure adequate data for model development. The final dataset included 22 birds (14,898 locations, 677 230 training steps per bird). For every observed step, 100 non-selected steps were generated with the R *amt* package (Signer *et al*. 2019), drawn from a uniform distribution ranging from 0 to 150% of the maximum observed step length (Rogers *et al*. 2025). All observed and non-selected steps were then annotated using the prepared environmental predictor layers. To obtain SSD predictions, iSSFs were fitted for every individual using the *amt* package, with environmental conditions at each step’s terminal point and a polynomial of step length as predictors (SI Table 1). These iSSFs were used to parametrize transition probability matrices, movements were restricted to the 99th percentile of observed step lengths (7214.5 m), and eigenvectors were computed to generate SSD surfaces for every individual (Fig. S1 XIV; Rogers *et al*. 2025). SDMs were fitted using the Maxent algorithm implemented in the *maxnet* R package (Phillips *et al*. 2017), based on the same environmental predictors (Table S1). GPS locations served as presence points, while 5,000 background points were randomly sampled from the modeled area. These models were used to project the probability of use across the surface (Fig. S1 XI). Resource selection functions (RSFs) were fitted for each individual using the same environmental predictors (Table S1), with 10 randomly generated pseudoabsence points per occurrence sampled within the minimum convex polygon of the combined locations (Signer *et al*. 2019), and used to project the predicted probability of use across the surface (Fig. S1 XII). Finally, the iSSFs used to generate the SSD projections were also used to create selection-only surfaces by predicting relative selection probabilities across the landscape, referenced to an arbitrary baseline cell (Fig. S1 XIII). All surfaces were kernelized and constrained to sum to 1. This procedure resulted in a kernelized SSD, SDM, RSF, and SSF space use surface per individual, which were subsequently averaged to generate population-level space use projections (Fig. S1). We focus on SDMs and SSDs in the main text, while results for RSF and SSF models are provided in the Supplementary Information for completeness (see Fig. S1 for schematic comparison; see Fig. S4 for population-level predictions).

#### Model evaluation

Individual-level prediction surfaces were evaluated using both the training data (75%) used during model fitting and the respective withheld testing data (25%; Fig. 4C “In-Sample” and “Out-of-Sample”). To assess model generalizability across individuals, each model was additionally tested against the combined data of all other individuals (Fig. 4C “Across Individuals”). Population-level predictions were evaluated by averaging individual prediction surfaces to create a mean population-level surface, which was then tested against the pooled set of testing data from all individuals (Fig. 4C “Population”). Further external validation was conducted using an independent dataset from annual aerial surveys carried out between 1950 and 2010 (Taylor *et al*. 2015). Those surveys targeted known whooping crane wintering areas and are assumed to cover the whole population’s wintering range. The full data set was used to determine the best estimate of the long-term distribution of crane space use and to minimize annual fluctuations (Fig. 4C “Independent”). Each evaluation was based on 100 bootstrap iterations, resampling the relevant testing data with replacement. The mean and 95% quantile intervals were computed to reduce sensitivity to variation in sampling composition. Since all predicted surfaces were kernelized to sum to one, the geometric mean of predicted intensity at known crane occurrence locations was used as the evaluation metric (following <u>Rogers *et al*.</u> <u>2025</u>; Fig. 4C).

The independent aerial survey data set was further used to explore correlations between model-predicted intensities and observed spatial population density. For visual comparisons, crane counts from aerial surveys were converted into a continuous density surface via kernel density estimation (KDE; Wand & Jones 1994; Fig. 4B, detailed description in Supporting Text S1). To quantify pixel-level spatial concordance between each model’s predicted surface and the empirical KDE map, two standard overlap metrics, Schoener’s D and Hellinger’s I (Warren *et al*. 2008), were computed. Because the aerial survey data have coarse spatial and temporal resolution and reflect aggregated use over large areas, model performance was further evaluated at the patch level. Habitat patches were defined as contiguous estuarine wetland areas as this is the dominant habitat type as indicated by model coefficients (Fig. S3) and visual alignment (Fig. 4A, Fig. S2). The distribution of resulting habitat patches is shown in Fig. S7. Observed density was calculated as crane counts per unit patch area, and mean predicted values were extracted for each patch. Because many small patches were unoccupied, we fit zero-inflated gamma models (*glmmTMB;* Brooks *et al*. 2017) with (i) a zero-inflation component for occupancy probability and (ii) a conditional component for density in occupied patches (SSD results shown in Fig. 4D; full comparison of all predictive methods in Fig. S8). Models were run at the population level using patch-aggregated predictors; attempts at mixed-effects models with random bird intercepts and slopes did not converge reliably.

#### Underlying mechanisms

To evaluate how key patch-level characteristics influence observed and predicted space use of Whooping Cranes, we tested the effects of habitat quality, patch size, and patch connectivity on both empirical density measures and model-based space use estimates. Habitat patches were defined as contiguous clusters of estuarine wetland pixels (as described in Section “*Model evaluation*“; patch distribution shown in Fig. S7). For each patch, we quantified three attributes: (i) proportional estuarine wetland cover as a measure of relative habitat quality, (ii) log-transformed patch size, and (iii) closeness centrality derived from patch-to-patch distance matrices as a proxy for connectivity (hereafter referred to as “patch closeness”). Observed space use was assessed using aerial survey-based density, defined as the number of cranes recorded during surveys from 1950–2010 per patch area. Because many patches were unused, we applied zero-inflated Gamma models to separately estimate the probability of use (zero-inflation component) and expected density within used patches (conditional component). Each patch attribute was tested in a separate univariate model. These analyses were intended to characterize whether observed and predicted space use showed similar broad relationships with each patch attribute, rather than to estimate their independent effects. Statistical significance was assessed via likelihood ratio tests, and predicted relationships were visualized along with 95% confidence intervals (Fig. 3A-C I). To test whether the same effects were reflected in model predictions, we extracted the average predicted use per patch from the SSD, SDM, RSF, and SSF projections. These predictions were based on population-level surfaces obtained by averaging across individuals, yielding expected use for a typical crane. For each method and patch attribute, univariate generalized linear models with a Gamma error distribution and log link were fitted using the *glmmTMB* package in R. Slope coefficients (β) and *p*-values were extracted to quantify effect strength and significance. All fitted relationships were visualized to illustrate predicted sensitivity to patch characteristics (SSD: Fig. 3A-C II, full comparisons in Fig. S9).

#### Future change projections

The Aransas region of the Texas Gulf Coast is expected to be significantly impacted by future sea level rise, which in turn could strongly affect crane habitats (Golden *et al*. 2022). To estimate how crane space use might shift under such sea level rise, the Marsh Migration dataset created by the NOAA Office for Coastal Management (https://coast.noaa.gov/slrdata/) was used for making future projections. The dataset models both loss of current marshes and creation of new marshes inland, depending on elevation, sea level rise, and natural sediment build-up (detailed method description available at https://coast.noaa.gov/data/digitalcoast/pdf/slr-marsh-migration-methods.pdf). The same reclassification approach used for current land cover maps (see Section “*Environmental predictors*“) was applied to the projected layers. The “Brackish/Transition Wetland” class appears only in the future layers and was assumed to be estuarine wetland to avoid model extrapolation. Global mean sea level is expected to rise between 0.43m (RCP2.6) and 0.84m (RCP8.5) by 2100 compared to 1986-2005 (Oppenheimer *et al*. 2019). Thus, three scenarios, 0.3m, 0.6m, and 0.9m, were considered in the analysis to examine forecasts of change in space use under low-, medium-, and high-rise conditions. SSD, SDM, RSF, and SSF models were projected onto each of the future environmental layers, and delta maps (current minus future predicted values) were calculated to assess changes in space use patterns. Future habitat disaggregation was assessed by calculating the change in average patch size and connectivity (quantified as closeness centrality as described previously) for future scenarios. Core areas of use were identified by determining the minimum area required to encompass 50% of total predicted space use, hereafter referred to as the “Min 50% Area.” Changes in space use concentration were quantified as the difference in Min 50% Area between current and future conditions.

## Results

### Individual movement captures drivers of population density

First, we examined whether consistent movement preferences are detectable at the scale of individual decisions. To do so, we leveraged 14,898 GPS locations from 22 individual Whooping Cranes in their wintering range in coastal Texas, USA (Fig. S2), to parameterize individual-level movement models. Our results reveal a clear and systematic structure in how individuals interact with their environment (Fig. 2). Step-selection functions show that cranes are more likely to make short rather than long movements between grid cells and to transition from lower- to higher-quality habitats rather than the reverse (Fig. 2A). When aggregating across all possible movement options that individuals can face in the landscape, these patterns remain evident: transitions are consistently biased towards nearby and higher-quality habitat (Fig. 2B).

**Figure 2:**
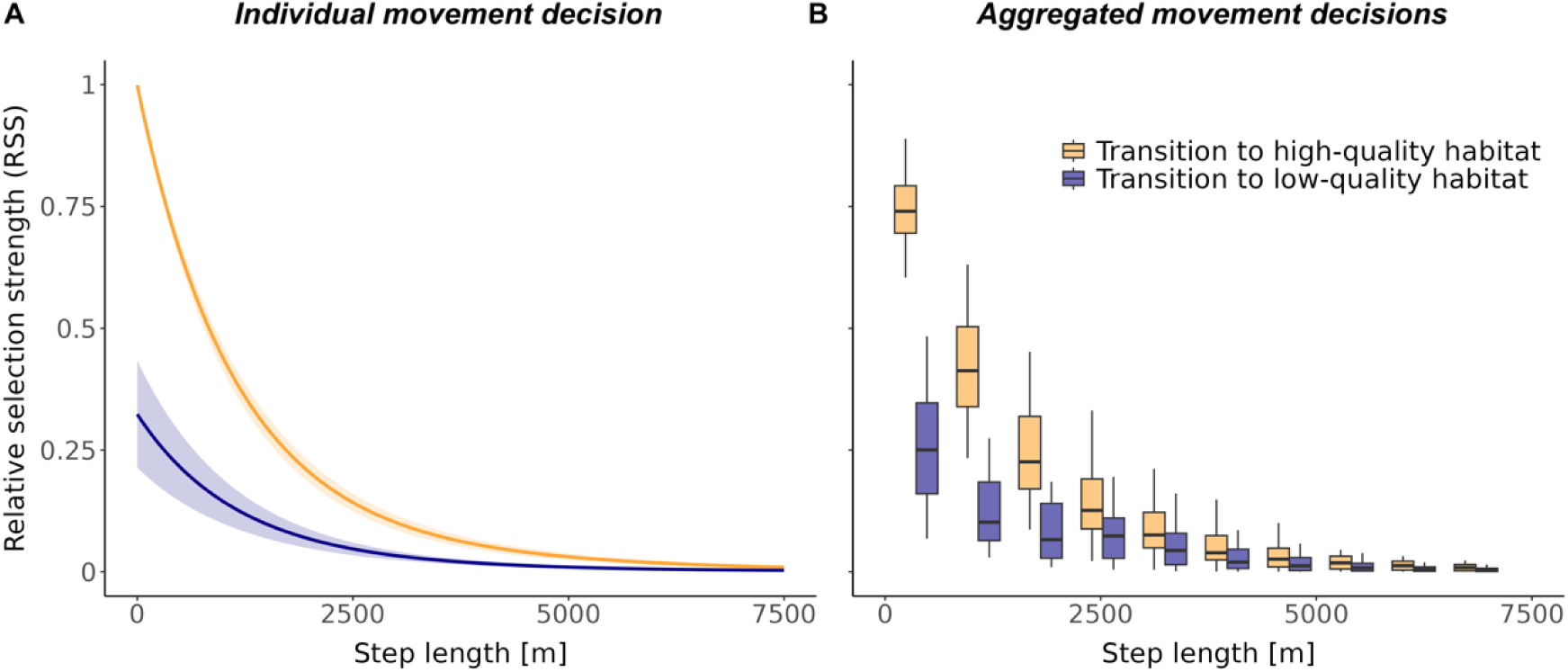
Local movement and selection generate landscape-scale connectivity biases. (A) Empirically estimated relative selection strength (RSS) for a single movement step as a function of step length when moving from a high-quality (estuarine wetland) origin cell into either another high-quality cell (orange) or a low-quality cell (purple). Lines show the mean SSF-predicted RSS across individuals (shaded bands indicate ± s.e), expressed relative to remaining in the same location (RSS = 1). Cranes preferentially make short movements into high-quality habitat, whereas long-distance moves and transitions into poorer habitat are progressively less likely. (B) Applying the same empirically derived step-selection rules across grid cells in the study region yields a landscape-scale network of potential transitions. Boxplots show the across-individual distribution of mean raw SSF-derived transition weights from high-quality origin cells to high- and low-quality destination cells across distance classes. The distance and selection biases underlying individual animal decisions give rise to general biases in how animals move through and use the landscape over time. Analytical details are provided in Supplementary Text S2.

These general biases toward short, high-quality movements (Fig. 2) imply that population density should accumulate in high-quality, expansive, and well-connected habitats. Indeed, independently collected, landscape-wide density estimates from aerial surveys confirm this qualitative expectation. Patch-level densities (cumulative counts per unit patch area), where patches are defined as contiguous estuarine wetland habitat, show significant positive associations with patch quality, size, and closeness when analyzed with zero-inflated gamma models (Fig. 3A-C).

**Figure 3:**
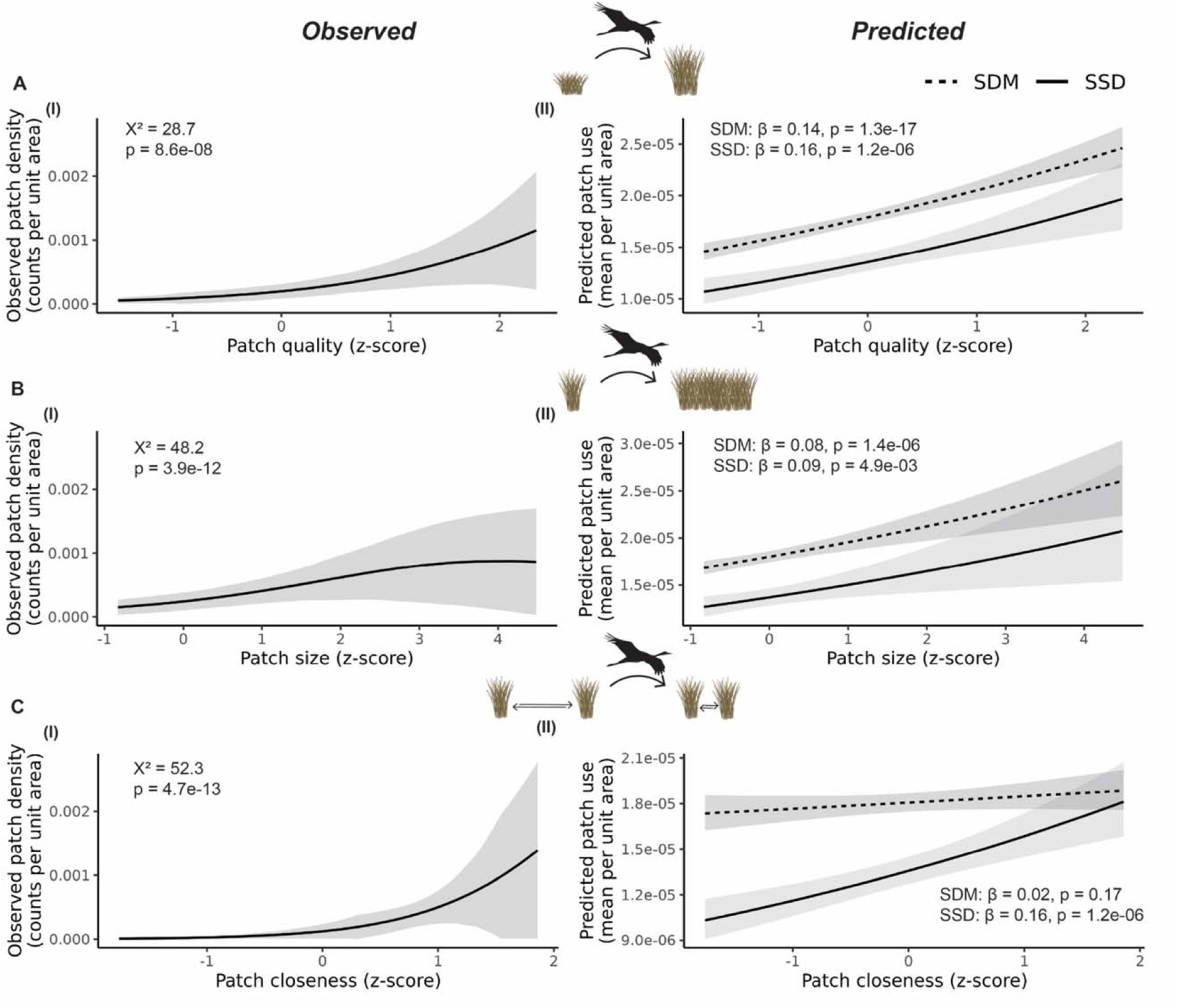
Movement links habitat configuration to spatial population density. Independently observed density estimates from aerial surveys, aggregated at the level of contiguous estuarine wetland habitat patches, increase with patch quality (A I), size (B I), and closeness (measured as closeness centrality; C I), indicating that large, high-quality, and well-connected habitats accumulate more individuals. Movement-informed SSDs reproduce these same relationships (A–C II), demonstrating that simple local movement and selection mechanisms are sufficient to generate the observed spatial structuring of populations across landscapes. In contrast, correlative SDMs capture the effect of patch quality (A II) and an apparent effect of patch size (B II; potentially through their collinearity; Fig. S10), but fail to capture the relationship with patch closeness (C II). χ² is the likelihood-ratio chi-squared statistic from comparing two nested zero-inflated Gamma GLMMs for the observed density data (null model vs. the model with the predictor in the zero-inflation part). β indicates the estimated slope from log link Gamma GLMs. All x-axes show standardized (z-score) values of the predictors.

We compared two models fitted to movement data in their ability to reflect these patterns: ergodic steady-state distributions (SSDs), which treat distributions as emerging from individual-level movement decisions, and correlative species distribution models (SDMs), which only use the occurrence information from the same data. Both models generate cell-level predictions that we aggregated to the patch level, enabling direct comparison with observed patch-level densities. While SDMs are expected to capture variation in habitat quality, mechanistic SSDs should additionally capture effects of habitat size and accessibility that arise through movement across the landscape. Consistent with this expectation, only movement-informed SSDs of crane space use exhibit significant positive associations with all three considered patch attributes: patch quality (Fig. 3A II), patch size (Fig. 3B II), and patch closeness (Fig. 3C II). Expectedly, SDMs yield positive relationships with patch quality (Fig. 3A II). But they capture an association with size (Fig. 3B II) only through the collinearity between size and quality (Fig. S10) and miss the role of connectivity (Fig. 3C II).

### Leveraging movement data for mapping population density

The patch-level relationships in Fig. 3 indicate that mechanistic SSDs capture all three considered landscape drivers of population density, whereas SDMs do not. We next asked whether this mechanistic consistency translates into improved spatial predictions of distribution and density. Although SDMs and SSDs are both fitted to the same movement data, they produce strikingly different landscape-scale predictions. SDMs identify high suitability wherever environmental conditions are favorable, assigning similar values to small, isolated habitat patches and to large, well-connected habitats (Fig. 4A). In contrast, SSDs generate spatially structured predictions that reflect not only habitat quality but also accessibility and configuration, distinguishing environmentally suitable habitat from habitat that is likely to contribute to realized space use (Fig. 4B).

**Figure 4:**
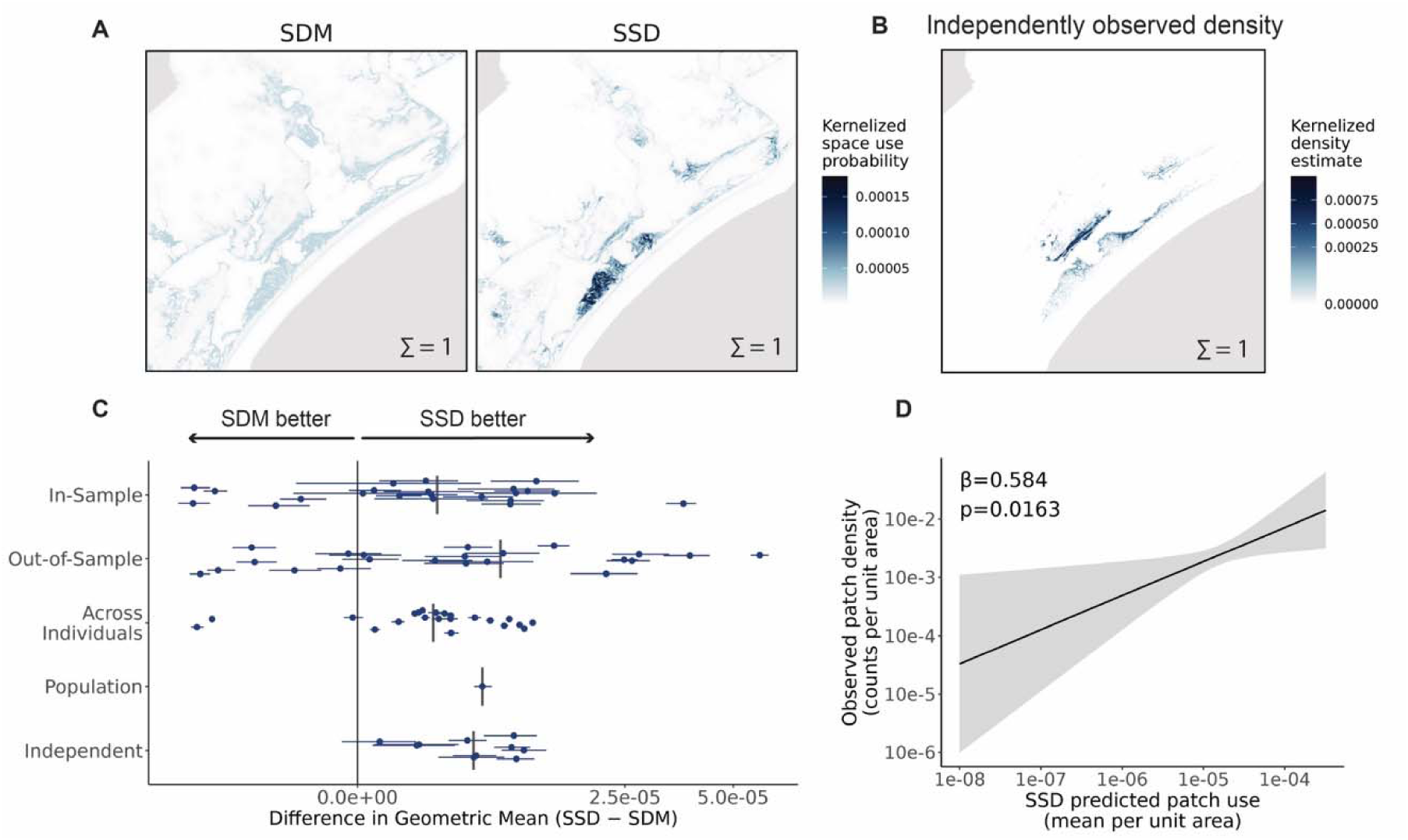
Models informed by movement mechanisms successfully predict spatial variation in population density. (A) Population-level predictions of crane space use from movement-informed SSDs and correlative SDMs, each normalized to sum to one (150 m resolution). SSDs concentrate predicted use in large, well-connected habitat complexes, whereas SDMs assign similarly high suitability to small and isolated patches. (B) Empirical kernel-density estimate of crane density from fully independent aerial surveys, showing strong spatial correspondence with SSD predictions. (C) Predictive performance of SSDs relative to SDMs, measured as the difference in geometric mean predicted values at observed locations (SSD − SDM). SSDs consistently assign higher predicted use to observed GPS and survey locations across in-sample (75 % of tracking data), out-of-sample (25 %), across-individual, population-level, and fully independent census validations, indicating improved discrimination between used and unused space. (D) SSD-predicted use is significantly associated with independently observed crane density (zero-inflated gamma: β = 0.58 ± 0.24, z = 2.40, *p* = 0.016), whereas analogous relationships for SDMs and other correlative approaches are not significant (full results in Fig. S8). Patch-level aggregation matches the spatial resolution at which aerial survey data are collected.

These differences are also evident quantitatively. SSDs consistently assign higher predicted use to observed GPS locations and independent survey locations than SDMs, indicating stronger discrimination between used and unused space (Fig. 4C). At the cell-level, SSDs show substantially greater agreement with observed spatial patterns of crane density, both visually and quantitatively (Schoener’s D: SSD = 0.23, SDM = 0.18; Hellinger’s I: SSD = 0.46, SDM = 0.39; Fig. 4A–C). To account for the coarse spatial and temporal resolution of the independent aerial surveys, we further evaluated predictions at the scale of habitat patches. Here again, only SSDs were significantly associated with observed patch-level densities conditional on occupancy (zero-inflated gamma; β = 0.58 ± 0.24, z = 2.40, p = 0.016; Fig. 4D), whereas SDMs showed no such relationship (Fig. S8).

### Projected response to environmental change

We finally assessed how incorporating movement-mediated connectivity in SSDs alters projections under global change, compared with traditional SDMs. Estuarine wetlands, identified as critical crane habitat (Fig. S3; Fig. S2), are projected to undergo substantial losses under sea level rise scenarios (Fig. 5A). In addition to overall habitat loss, these changes cause a change in spatial habitat cohesion, with once-contiguous patches becoming increasingly fragmented and isolated. Under increasing sea-level rise, both the average size and connectivity (here closeness centrality) of suitable habitat patches decline by up to 80% (Fig. 5C).

**Figure 5:**
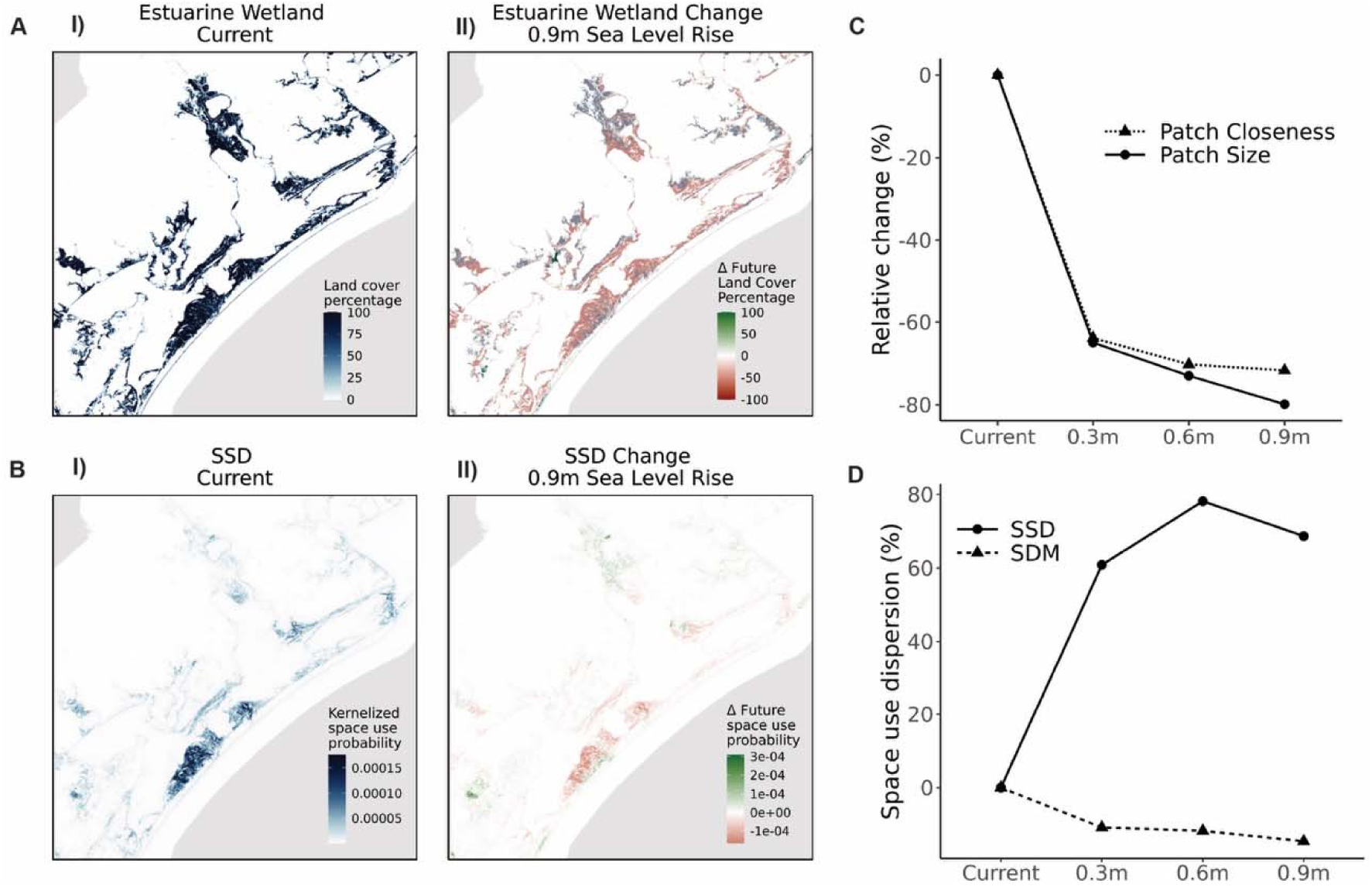
Movement mediates how habitat loss reorganizes populations. (A) Current estuarine wetland cover and the same layer overlaid with projected changes under 0.9 m sea-level rise (SLR), showing widespread loss of habitat and increasing fragmentation of remaining wetlands. (B) Corresponding current SSD-predicted space use and projected change under SLR, revealing strong contraction of use in lost habitat and intensified concentration in the remaining large, well-connected patches. (C) Patch-level habitat structure under increasing SLR scenarios, showing steep declines in both patch size and patch closeness (quantified as closeness centrality) as wetlands are fragmented. (D) Predicted space use dispersion (quantified as the percent change in the minimum area required to contain 50% of the predicted population-level space-use probability) under SLR. Movement-informed SSDs predict increasing dispersion and reorganization of space use as habitat fragments, whereas correlative SDMs show little or no response. This highlights SSD’s ability to capture spatial reorganization in response to landscape change. For full comparison of all methods and sea level rise scenarios, see Fig. S11.

SSD models accordingly project pronounced shifts in crane space use and population redistributions. Space use declines sharply in large, previously well-connected coastal patches lost to inundation, while increasing further inland in smaller, more isolated fragments that persist (Fig. 5A-B). This results in a marked increase in the spatial dispersion of space use (measured as the minimum area required to encompass 50% of predicted space use), specifically an up to 80% expansion of core use area, as cranes are forced to rely on scattered, lower-quality habitat in the future (Fig. 5D). In SDMs, the effects of future inundation are only captured through changes in local environmental conditions, without explicitly accounting for changes in patch size or connectivity, leading to predictions of much lower levels of change (Fig. 5D; Fig. S11).

## Discussion

The spatial structure of animal populations is determined not only by where suitable habitat exists, but also by how individuals access it as the move across landscapes. Here we show that individual-level movement dynamics can generate emergent population-level distributions and densities by jointly shaping habitat selection and accessibility. Using GPS tracking data, we estimated such individual-level decision processes empirically and projected them across a complex landscape. As animals bias their movements toward nearby, high-quality habitat (Fig. 2), large, connected, or centrally located areas accumulate use, while equally suitable yet isolated patches remain underused (Fig. 3A-C I). Derived space use based on local movement mechanisms captured these crucial effects of habitat configuration (Fig. 3A-C II and, despite being parameterized from a relatively small sample of individuals, captured key features of independently collected, landscape-wide population density data (Fig. 4). Building on this present-day agreement, projections under sea-level rise forecast severe population redistribution as habitat is lost and spatially reconfigured – changes that conventional correlative approaches largely fail to capture (Fig. 5).

Our results highlight how animal movement information can empirically integrate three complementary dimensions of population distributions: habitat suitability from niche theory, habitat accessibility from connectivity science, and the resulting spatial variation in population use and density emphasized by metapopulation ecology. By linking these dimensions through observed movement decisions, movement-informed models provide a mechanistic explanation for why static, correlative models can misinterpret where animals occur (Van Moorter *et al*. 2023; Velazco *et al*. 2020). SDMs usually project the environmental niche onto geographic space under the assumption of uniform accessibility, neglecting distances, barriers, and corridors that constrain movement in the landscape (Rogers *et al*. 2025; Van Moorter *et al*. 2023). In doing so, they effectively flatten the accessibility gradients that underlie population density (Pulliam 2000; Soberón & Peterson 2005; Fig. 2, Fig. 3). As a result, suitable but isolated areas may appear highly suitable even where accessibility constraints reduce their functional value to the population (Fig. 4, Fig. S5). Here leveraging empirical movement mechanism provides a clear advantage: accessibility gradients are inherently accounted for in predictions and observed variation in use of habitat is recovered (Fig. 3) resulting in improved alignment with independent population density data (Fig. 4). More broadly, acknowledging accessibility as a key driver of space use shifts distribution modelling from mapping potential habitat to inferring realized use. Identifying where suitable habitat is also functionally accessible can help reveal important use hotspots and direct conservation toward areas that sustain and concentrate movement trajectories (Van Moorter *et al*. 2023). Because movement is a fundamental process shaping how mobile organisms encounter and use heterogeneous landscapes across ecosystems, this principle may have broad relevance across taxa and regions globally.

Habitat loss is a multifaceted problem, composed of reductions in habitat extent, deterioration of habitat quality, and increased fragmentation of remaining habitats (Haddad *et al*. 2015; Hansen *et al*. 2013). For mobile species, our work suggests that habitat fragmentation may exacerbate biodiversity loss far beyond the absolute reduction of suitable habitat (Fletcher *et al*. 2018; Haddad *et al*. 2015). By accounting for the spatial arrangement of habitat, movement-informed projections of space use forecast a major population redistribution and dispersion under future landscape change. In contrast, traditional approaches only capture changes of habitat abundance in future suitability estimates (Fig. 5). Although this is a known issue in traditional occurrence-based frameworks like SDMs (Carvajal *et al*. 2018; Halstead *et al*. 2019) such prediction frameworks remain widely used in predicting current and future distributions of highly mobile organisms (Lan *et al*. 2025; Porto *et al*. 2024; Sierra-Morales *et al*. 2021). As a result, future projections of suitable habitat under anthropogenic change may be over-optimistic in increasingly fragmented landscapes. Our results demonstrate the negative implications of fragmentation-induced reductions in the remaining habitat functionality for mobile organisms (Van Moorter *et al*. 2023). As the wealth of animal biologging data continues to grow (Jetz *et al*. 2022; Kays *et al*. 2015; Kays & Wikelski 2023), leveraging the potential of the movement mechanisms inherently contained in this data is critical to enhance conservation efforts and to accurately anticipate biodiversity change.

Together, our results indicate that much of the spatial structure of populations can be explained and predicted by the interaction of empirically estimated movement mechanisms with landscape structure. Whereas metapopulation models (Hanski 1999, 1998) describe how patch networks influence persistence through stochastic extinction and recolonization, our movement-based framework explains how spatial structure emerges even within a single, continuously occupied population. While other ecological drivers, such as biotic interactions (Wisz *et al*. 2013), memory (Kim *et al*. 2024), or disease (Dougherty *et al*. 2018), can further shape spatial population structure, our findings suggest that movement and selection alone can capture the dominant spatial gradients of observed space use. Moreover, our estimates were based on a relatively small sample of individuals. Because predictions are derived from empirical, individual-level mechanisms rather than population-level correlations, movement mechanisms offer more natural extensions to other individuals or spatial extents where mechanisms can be assumed to remain valid. While the expression of movement mechanisms may differ among individuals or demographic groups such as age or sex (Carlson *et al*. 2021; Lu *et al*. 2025), may not capture rare behaviors, or vary through time (Raynor *et al*. 2017; Tsalyuk *et al*. 2019) and under novel environmental conditions, mechanistic movement-informed predictions offer important insights into the rules shaping population spatial distributions.

Under ever-growing anthropogenic change, understanding population responses to landscape alterations is critical to conserve biodiversity and its many functions and services (Powers & Jetz 2019; Visconti *et al*. 2016). While metapopulation models have long addressed how habitat quality and structure jointly shape population persistence, their empirical application typically requires extensive population monitoring across large spatiotemporal extents. For many species, such data remain difficult to obtain, limiting the scalable use of these frameworks for predicting continuous population distributions and densities. Conservation planning therefore often relies on correlative SDM approaches, which address different ecological questions and are comparatively data-efficient, but do not inherently represent habitat geometry or movement-mediated accessibility. Our work suggests that empirically estimated, individual-level movement mechanisms offer an effective and scalable alternative. Individuals are the building blocks of populations, and how these individuals navigate complex landscapes offers insights into the drivers of animal movement (Avgar *et al*. 2016). Beyond describing movement behavior, empirically derived movement mechanisms can be used to understand individual-level space use (Avgar *et al*. 2016; Michelot *et al*. 2019; Potts & Börger 2023; Rogers *et al*. 2025; Van Moorter *et al*. 2023). Our work suggests that these individual-level mechanisms scale up, allowing patterns inferred from few individuals to describe the space use of an entire population. More work is could help refine our understanding of how individual-level mechanisms inform population-level outcomes (Carlson *et al*. 2021; Lu *et al*. 2025), including how density dependence and social interactions may modify movement decisions and realized space use. Critically, we show that ignoring these individual-level mechanisms, particularly movement in mobile species, may underestimate the impacts of anthropogenic change. Movement-informed models, applied to the rapidly growing body of GPS-tracking and satellite data, therefore represent a promising avenue to mechanistically informed yet scalable monitoring and decision-support for biodiversity conservation (Jetz *et al*. 2022). In particular, they are critical for understanding, maintaining, and restoring landscape connectivity, and thus the functional integrity and resilience of ecosystems in an increasingly fragmented world.

## Acknowledgments

We thank Niklaus Zimmermann for his advice and feedback throughout the creation of the manuscript. Thanks also to the Spatial Modeling Group at the Yale Center for Biodiversity and Global Change, including Shubhi Sharma, Richard Li, and Jeremy Cohen, for their feedback on figures and versions of this manuscript. Any use of trade, firm, or product names is for descriptive purposes only and does not imply endorsement by the U.S. Government.

## References

Avgar, T., Lele, S.R., Keim, J.L. & Boyce, M.S. (2017). Relative Selection Strength: Quantifying effect size in habitat- and step-selection inference. Ecology and Evolution, 7, 5322–5330.

Avgar, T., Potts, J.R., Lewis, M.A. & Boyce, M.S. (2016). Integrated step selection analysis: bridging the gap between resource selection and animal movement. Methods in Ecology and Evolution, 7, 619–630.

Bascompte, J. & Solé, R.V. (1996). Habitat Fragmentation and Extinction Thresholds in Spatially Explicit Models. Journal of Animal Ecology, 65, 465–473.

Brodie, J.F., Gonzalez, A., Mohd-Azlan, J., Nelson, C.R., Tabor, G., Vasudev, D., et al. (2025). A well-connected Earth: The science and conservation of organismal movement. Science, 388, eadn2225.

Brooks, M.E., Kristensen, K., van Benthem, K.J., Magnusson, A., Berg, C.W., Nielsen, A., et al. (2017). glmmTMB Balances Speed and Flexibility Among Packages for Zero-inflated Generalized Linear Mixed Modeling. The R Journal, 9, 378–400.

Carlson, B.S., Rotics, S., Nathan, R., Wikelski, M. & Jetz, W. (2021). Individual environmental niches in mobile organisms. Nature Communications, 12, 4572.

Carvajal, M.A., Alaniz, A.J., Smith-Ramírez, C. & Sieving, K.E. (2018). Assessing habitat loss and fragmentation and their effects on population viability of forest specialist birds: Linking biogeographical and population approaches. Diversity and Distributions, 24, 820–830.

Dougherty, E., Seidel, D., Carlson, C., Spiegel, O. & Getz, W. (2018). Going through the motions: Incorporating movement analyses into disease research. Ecology Letters, 21.

Elith, J. & Leathwick, J.R. (2009). Species Distribution Models: Ecological Explanation and Prediction Across Space and Time. Annual Review of Ecology, Evolution, and Systematics.

Ellison-Neary, N., Oppenheimer, S., Dehnen, T., Hatchwell, B.J. & Street, G.M. (2026). An intuitive method to calculate the utilization distribution of an animal from step-selection analysis. Methods in Ecology and Evolution, 17, 2175–2188.

Fahrig, L. (2017). Ecological Responses to Habitat Fragmentation Per Se. Annual Review of Ecology, Evolution, and Systematics.

Fletcher Jr., R.J., Sefair, J.A., Wang, C., Poli, C.L., Smith, T.A.H., Bruna, E.M., et al. (2019). Towards a unified framework for connectivity that disentangles movement and mortality in space and time. Ecology Letters, 22, 1680–1689.

Fletcher, R.J., Didham, R.K., Banks-Leite, C., Barlow, J., Ewers, R.M., Rosindell, J., et al. (2018). Is habitat fragmentation good for biodiversity? Biological Conservation, 226, 9–15.

Golden, K.E., Hemingway, B.L., Frazier, A.E., Scholtz, R., Harrell, W., Davis, C.A., et al. (2022). Spatial and temporal predictions of whooping crane (Grus americana) habitat along the US Gulf Coast. Conservation Science and Practice, 4, e12696.

Guisan, A., Tingley, R., Baumgartner, J.B., Naujokaitis-Lewis, I., Sutcliffe, P.R., Tulloch, A.I.T., et al. (2013). Predicting species distributions for conservation decisions. Ecology Letters, 16, 1424–1435.

Guisan, A. & Zimmermann, N.E. (2000). Predictive habitat distribution models in ecology. Ecological Modelling, 135, 147–186.

Haddad, N.M., Brudvig, L.A., Clobert, J., Davies, K.F., Gonzalez, A., Holt, R.D., et al. (2015). Habitat fragmentation and its lasting impact on Earth’s ecosystems. Science advances, 1, e1500052.

Halstead, K.E., Alexander, J.D., Hadley, A.S., Stephens, J.L., Yang, Z. & Betts, M.G. (2019). Using a species-centered approach to predict bird community responses to habitat fragmentation. Landscape Ecology, 34, 1919–1935.

Hansen, M.C., Potapov, P.V., Moore, R., Hancher, M., Turubanova, S.A., Tyukavina, A., et al. (2013). High-Resolution Global Maps of 21st-Century Forest Cover Change. Science, 342, 850–853.

Hanski, llkka. (1999). Metapopulation Ecology. Oxford University Press. Hanski, I. (1998). Metapopulation dynamics. Nature, 396, 41–49.

Hanski, I. (2011). Habitat Loss, the Dynamics of Biodiversity, and a Perspective on Conservation. AMBIO, 40, 248–255.

Jetz, W., Tertitski, G., Kays, R., Mueller, U., Wikelski, M., Åkesson, S., et al. (2022). Biological Earth observation with animal sensors. Trends in Ecology & Evolution, 37, 293–298.

Kays, R., Crofoot, M.C., Jetz, W. & Wikelski, M. (2015). Terrestrial animal tracking as an eye on life and planet. Science, 348, aaa2478.

Kays, R. & Wikelski, M. (2023). The Internet of Animals: what it is, what it could be. Trends in Ecology & Evolution, 38, 859–869.

Kim, D., Thompson, P.R., Wolfson, D.W., Merkle, J.A., Oliveira-Santos, L.G.R., Forester, J.D., et al. (2024). Identifying signals of memory from observations of animal movements. Movement Ecology, 12, 72.

Lan, Y., Wu, X., Xu, M., Li, K., Huan, Y., Zhou, G., et al. (2025). High-resolution global distribution projections of 10 rodent genera under diverse SSP-RCP scenarios, 2021–2100. Scientific Data, 12, 1467.

Lehnen, S.E., Sesnie, S.E., Butler, M.J., Pearse, A.T. & Metzger, K.L. (2024). Management implications of habitat selection by whooping cranes (Grus americana) on the Texas coast. Ecosphere, 15, e4820.

Lu, M., Yanco, S.W., Carlson, B.S., Winner, K., Cohen, J.M., Ellis-Soto, D., et al. (2025). A theoretical framework for scaling ecological niches from individuals to species. Proceedings of the National Academy of Sciences, 122, e2425582122.

McGarigal, K. (1995). FRAGSTATS: spatial pattern analysis program for quantifying landscape structure. US Department of Agriculture, Forest Service, Pacific Northwest Research Station.

Metzger, K.L., Lehnen, S.E., Sesnie, S.E., Butler, M.J., Pearse, A.T. & Harris, G. (2020). Identifying sustainable winter habitat for whooping cranes. Journal for Nature Conservation, 57, 125892.

Michelot, T., Blackwell, P.G. & Matthiopoulos, J. (2019). Linking resource selection and step selection models for habitat preferences in animals. Ecology, 100, e02452.

Moorcroft, P.R., Lewis, M.A. & Crabtree, R.L. (2006). Mechanistic home range models capture spatial patterns and dynamics of coyote territories in Yellowstone. Proceedings of the Royal Society B: Biological Sciences, 273, 1651–1659.

Mueller, T. & Fagan, W.F. (2008). Search and navigation in dynamic environments – from individual behaviors to population distributions. Oikos, 117, 654–664.

Nathan, R., Getz, W.M., Revilla, E., Holyoak, M., Kadmon, R., Saltz, D., et al. (2008). A movement ecology paradigm for unifying organismal movement research. Proceedings of the National Academy of Sciences, 105, 19052–19059.

National Oceanic and Atmospheric Administration, Office for Coastal Management. (2016). “2016 NOAA C-CAP Regional Land Cover” Coastal Change Analysis Program (C-CAP) Regional Land Cover.

Oppenheimer, M., Glavovic, B.C., Hinkel, J., van de Wal, R., Magnan, A.K., Abd-Elgawad, A., et al. (2019). Sea Level Rise and Implications for Low-Lying Islands, Coasts and Communities. In: IPCC Special Report on the Ocean and Cryosphere in a Changing Climate (eds.

Pörtner, H.-O., Roberts, D.C., Masson-Delmotte, V., Zhai, P., Tignor, M., Poloczanska, E., et al.). Cambridge University Press, Cambridge, UK and New York, NY, USA, pp. 321–445.

Pearse, A.T., Brandt, D.A., Baasch, D.M., Bidwell, M.T., Conkin, J.A., Harner, M.J., et al. (2020). Location data for whooping cranes of the Aransas-Wood Buffalo Population, 2009-2018.

Phillips, S.J., Anderson, R.P., Dudík, M., Schapire, R.E. & Blair, M.E. (2017). Opening the black box: an open-source release of Maxent. Ecography, 40, 887–893.

Porto, L.M.V., Bennett, D., Maestri, R. & Etienne, R.S. (2024). Canids in a changing climate: predicting range shifts and evolutionary rescue under distinct future scenarios. Biological Journal of the Linnean Society, 143, blae094.

Potts, J.R. & Börger, L. (2023). How to scale up from animal movement decisions to spatiotemporal patterns: An approach via step selection. Journal of Animal Ecology, 92, 16–29.

Powers, R.P. & Jetz, W. (2019). Global habitat loss and extinction risk of terrestrial vertebrates under future land-use-change scenarios. Nature climate change, 9, 323–329.

Pressey, R.L., Cabeza, M., Watts, M.E., Cowling, R.M. & Wilson, K.A. (2007). Conservation planning in a changing world. Trends Ecol Evol, 22, 583–592.

Pulliam, H.R. (2000). On the relationship between niche and distribution. Ecology Letters, 3, 349–361.

Raynor, E.J., Beyer, H.L., Briggs, J.M. & Joern, A. (2017). Complex variation in habitat selection strategies among individuals driven by extrinsic factors. Ecology and Evolution, 7, 1802–1822.

Rogers, W., Yanco, S. & Jetz, W. (2025). Choices to Landscapes: Mechanisms of Animal Movement Scale to Landscape Patterns of Space Use. Ecology Letters, 28, e70279.

Sierra-Morales, P., Rojas-Soto, O., Ríos-Muñoz, C.A., Ochoa-Ochoa, L.M., Flores-Rodríguez, P. & Almazán-Núñez, R.C. (2021). Climate change projections suggest severe decreases in the geographic ranges of bird species restricted to Mexican humid mountain forests. Global Ecology and Conservation, 30, e01794.

Signer, J., Fieberg, J. & Avgar, T. (2019). Animal movement tools (amt): R package for managing tracking data and conducting habitat selection analyses. Ecology and Evolution, 9, 880–890.

Soberón, J. (2007). Grinnellian and Eltonian niches and geographic distributions of species. Ecology Letters, 10, 1115–1123.

Soberón, J. & Peterson, A.T. (2005). Interpretation of Models of Fundamental Ecological Niches and Species’ Distributional Areas. Biodiversity Informatics, 2.

Swift, T.L. & Hannon, S.J. (2010). Critical thresholds associated with habitat loss: a review of the concepts, evidence, and applications. Biological Reviews, 85, 35–53.

Taylor, L.N., Ketzler, L.P., Rousseau, D., Strobel, B.N., Metzger, K.L. & Butler, M.J. (2015). Observations of whooping cranes during winter aerial surveys: 1950-2011. Aransas National Wildlife Refuge, U.S. Fish and Wildlife Service, Austwell, Texas, USA.

Thomas, J.A., Bourn, N.A.D., Clarke, R.T., Stewart, K.E., Simcox, D.J., Pearman, G.S., et al. (2001). The quality and isolation of habitat patches both determine where butterflies persist in fragmented landscapes. Proceedings of the Royal Society of London. Series B: Biological Sciences, 268, 1791–1796.

Tsalyuk, M., Kilian, W., Reineking, B. & Getz, W.M. (2019). Temporal variation in resource selection of African elephants follows long-term variability in resource availability. Ecological Monographs, 89, e01348.

Urban, D. & Keitt, T. (2001). LANDSCAPE CONNECTIVITY: A GRAPH-THEORETIC PERSPECTIVE. Ecology, 82, 1205–1218.

Van Moorter, B., Kivimäki, I., Panzacchi, M., Saura, S., Brandão Niebuhr, B., Strand, O., et al. (2023). Habitat functionality: Integrating environmental and geographic space in niche modeling for conservation planning. Ecology, 104, e4105.

Velazco, S.J.E., Ribeiro, B.R., Laureto, L.M.O. & Júnior, P.D.M. (2020). Overprediction of species distribution models in conservation planning: A still neglected issue with strong effects. Biological Conservation, 252, 108822.

Venter, O., Sanderson, E.W., Magrach, A., Allan, J.R., Beher, J., Jones, K.R., et al. (2016). Sixteen years of change in the global terrestrial human footprint and implications for biodiversity conservation. Nature Communications, 7, 12558.

Visconti, P., Bakkenes, M., Baisero, D., Brooks, T., Butchart, S.H.M., Joppa, L., et al. (2016). Projecting Global Biodiversity Indicators under Future Development Scenarios. Conservation Letters, 9, 5–13.

Wand, M.P. & Jones, M.C. (1994). Kernel smoothing. CRC press.

Warren, D.L., Glor, R.E. & Turelli, M. (2008). ENVIRONMENTAL NICHE EQUIVALENCY VERSUS CONSERVATISM: QUANTITATIVE APPROACHES TO NICHE EVOLUTION. Evolution, 62, 2868–2883.

Wisz, M.S., Pottier, J., Kissling, W.D., Pellissier, L., Lenoir, J., Damgaard, C.F., et al. (2013). The role of biotic interactions in shaping distributions and realised assemblages of species: implications for species distribution modelling. Biological Reviews, 88, 15–30.

