## Supplementary Information for "Movement mechanisms explain current animal density patterns and predict their reorganization under global change"

**This PDF file includes:**

Supporting Text S1 to S2

Figures S1 to S12

Table S1

SI References

Supporting Text S1: Description of KDE surface creation

KDE is a non-parametric smoothing method that turns discrete point observations into a continuous surface by placing a weighted Gaussian kernel around each observation (Wand & Jones 1994). Each observation was weighted by the reported number of birds to ensure proportional representation in the resulting surface. The smoothing parameter controls the spread of each kernel and influences the level of detail in the resulting density surface. To determine an optimal smoothing scale, the data-driven bandwidth selection method, *bw.diggle* from the *spatstat* package, was used (Baddeley *et al.* 2015). The resulting density map was then resampled to align with the model prediction grids and kernelized to sum to 1 across the region (Fig. 4B).

Supporting Text S2: Computation of movement selection patterns in Fig. 2

Panel A: We generated a sequence of 7,500 hypothetical step lengths (1–7,500 in the model’s step-length units, corresponding to meters in the projected coordinate system) and used the fitted individual step-selection function (iSSF) models to predict the relative selection strength (RSS) for two contrasting destination environments: (i) movement from a high-quality origin cell into a high-quality destination cell, and (ii) movement from a high-quality origin cell into a low-quality destination cell. High-quality habitat was defined as a cell with 100% *Estuarine Wetland* cover, whereas low-quality habitat was defined as a cell with 100% *Upland* cover. To isolate the effect of step length and destination habitat, the shoreline-distance covariate was fixed at its landscape-wide mean. For each bird, linear predictors were obtained from the fitted iSSF for each hypothetical step and exponentiated relative to a reference step of zero length (i.e. remaining in the same location), yielding $RSS\left( d \right)=exp(\eta(d)-\eta(0))$ where $\eta(d)$ denotes the linear predictor for a step of length d. An RSS of 1 indicates that moving a given distance is equally likely as remaining in place. Mean RSS (± s.e.) across individuals was plotted to illustrate how cranes preferentially make short movements and transitions into high-quality habitat, whereas long steps and moves into low-quality habitat are comparatively disfavoured (Fig. 2A).

Panel B: To assess how the iSSF’s step-selection biases vary across the landscape context represented in the rasterized habitat layers, we recomputed the SSF-based transition matrices used for the SSD analysis without row normalization, thereby retaining the raw step-selection weights $w\left( i\to j \right)=exp(\eta_{i}-\eta_{j})$, rather than normalized transition probabilities. Row normalization was omitted because the subsequent filtering steps remove subsets of destinations from each origin, which would invalidate row-standardized probabilities. We restricted origins to high-quality habitat cells (≥ 90% Estuarine Wetland cover) and retained only destinations whose standardized shoreline-distance covariate matched that of the origin and that were classified as either high-quality (≥ 90% Estuarine Wetland) or low-quality (≤ 10% Estuarine Wetland). These threshold-based classes provide a landscape-based analogue of the idealized habitat contrasts used in Panel A. All valid transitions were grouped into ten equally spaced distance bins spanning the full range of observed neighbour distances and separated by transition type (high-quality → high-quality vs. high-quality → low-quality). For each origin cell, valid outgoing transitions were grouped into ten equally spaced distance bins spanning the full range of neighbor distances and separated by transition type (high-quality → high-quality vs. high-quality → low-quality). Within each origin, we computed the mean transition weight in each distance bin and transition type, and then averaged these per-origin means to obtain bird-specific summaries. These bird-level summaries were then aggregated across individuals for plotting. The resulting curves summarize how relative transition weight varies with step length for movements from high-quality habitat into high- versus low-quality destinations, showing stronger short-distance transitions into high-quality habitat and comparatively weaker long-distance or downward-quality transitions (Fig. 2B).

Figures

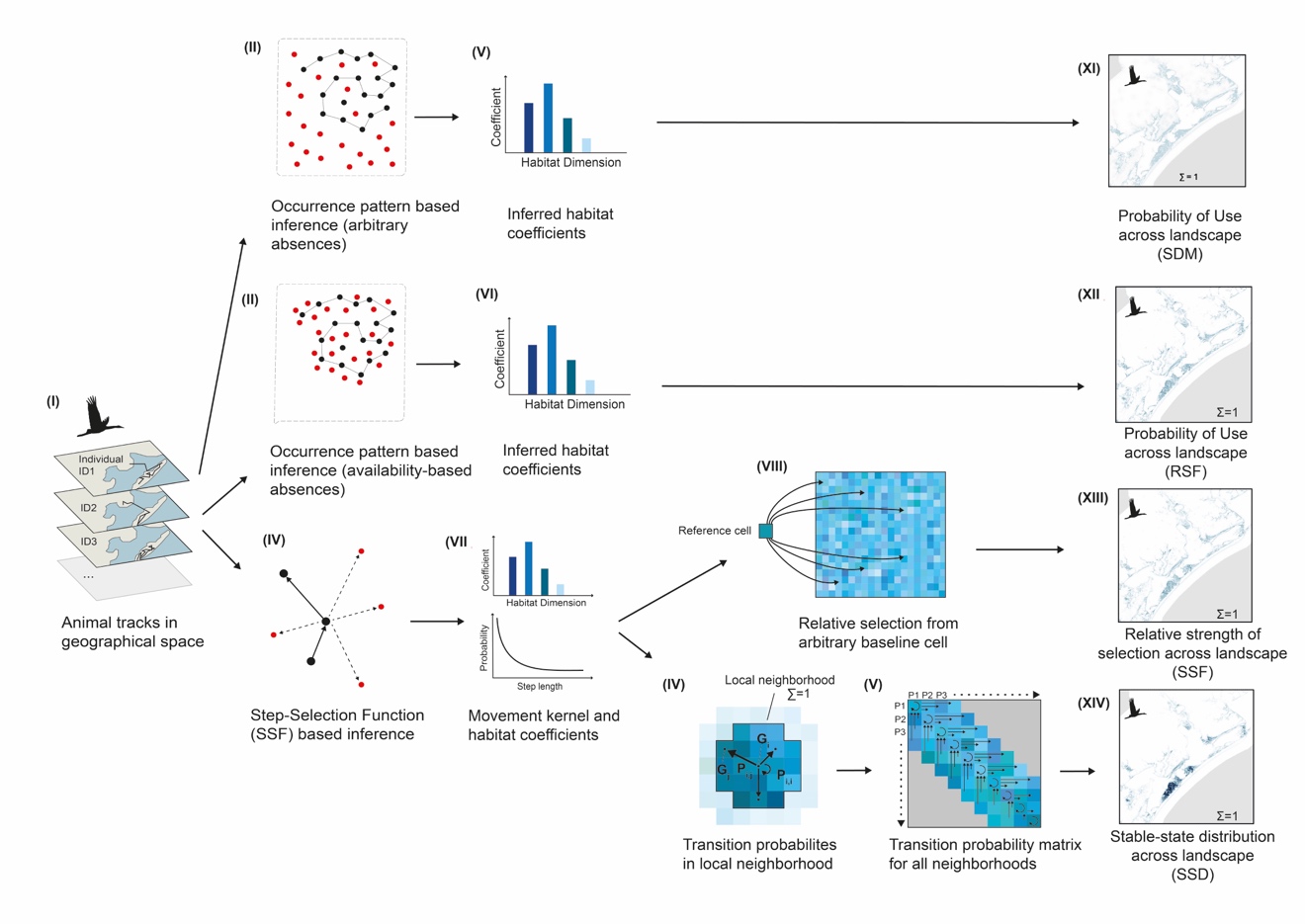

Fig. S1: Schematic overview of space use modelling approaches

First, animal movement data (I) can be used to generate spatial predictions of habitat use through four common modeling approaches. First, recorded locations can be treated as presence-only data and used in traditional species distribution models (SDMs) such as Maxent, which contrast observed locations with arbitrary background points (II). These models estimate habitat coefficients in environmental space (V) and produce relative habitat suitability surfaces (XI) which can be normalized for comparability.

Second, movement data can be analyzed using resource selection functions (RSFs), which contrast observed locations with availability-based absences sampled from each individual's accessible environment (III). RSFs similarly estimate habitat coefficients (VI) and yield a probability of use surface (XII) that can also be normalized.

Third, movement paths can be interpreted as transitions between used and available steps (IV), enabling inference via step selection functions (SSFs). Integrated SSFs (iSSFs) estimate both habitat coefficients and a movement kernel (VII), and can be used to predict the relative strength of selection for each location in comparison to an arbitrary reference cell (VIII). The resulting spatial surface (XIII) is structurally similar to RSF outputs but grounded in mechanistic, movement-informed coefficients.

Finally, iSSF parameters can be used to estimate transition probabilities between each cell and its local neighbors (IX), producing a complete transition probability matrix across the landscape (X). The first principal eigenvector of this matrix defines the stable-state distribution (SSD) of the system (XIV), which represents the long-term equilibrium distribution of space use implied by repeated local movements. Unlike SDM, RSF, and SSF projections, SSD explicitly incorporates both local selection and movement dynamics.

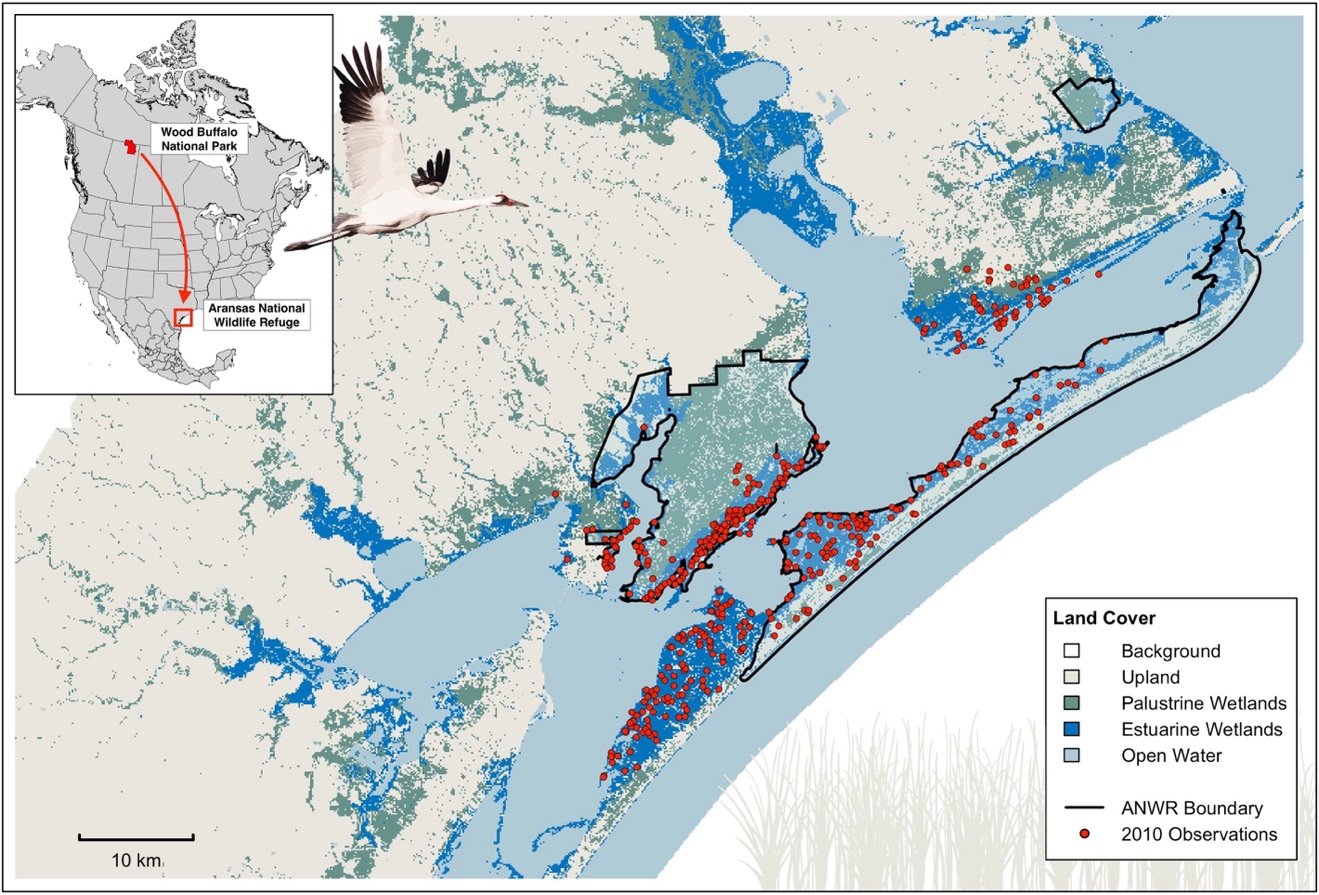

Fig. S2: Study area and landcover distribution

Wintering grounds of the last self-sustaining population of whooping cranes (*Grus americana*) along the Texas Gulf Coast. The study area encompasses the landscape surrounding the Aransas National Wildlife Refuge (ANWR), outlined in black. Aggregated landcover layers are shown to illustrate habitat heterogeneity. Red points represent whooping crane locations recorded during the final year (2010) of a long-term aerial survey campaign (Taylor *et al.* 2015). Shape files were rerieved from (Commission for Environmental Cooperation (CEC) *et al.* 2022; UNEP-WCMC 2025). Land cover was derived form NOAA Marsh Migration dataset from 2016 (<https://coast.noaa.gov/slrdata/>) as described in the method section of the main manuscript.

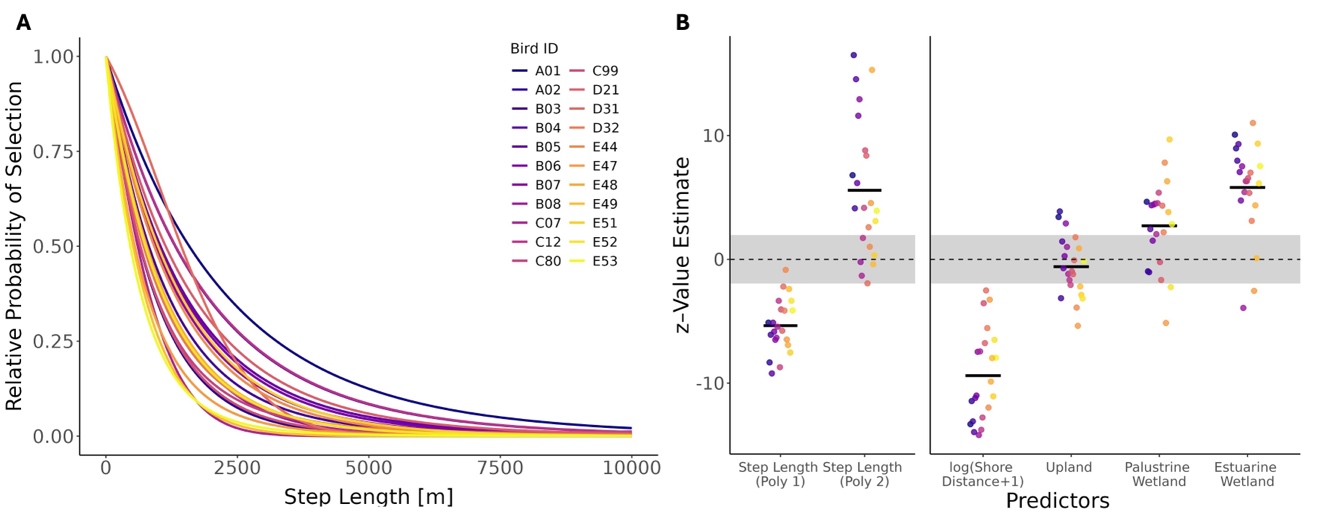

Fig. S3: SSF/SSD habitat coefficients and movement kernels

**(A)** iSSF functions as used in this analysis include a movement term that captures the likelihood of making movements of different step lengths. This plot shows the relative probability of taking a step from a cell to another cell with the same habitat conditions under varying step lengths. The maximum shown here is 10.000 m, the maximal step length considered in the transition probability matrices during the SSD generation was 7214.5 m (99th percentile of step length distribution). Colors indicate models for different individuals in the population. Curves show similar forms, but variations exist between individuals as they are derived from different step length distributions based on the movement behavior of each animal. **(B)** Z-values of the habitat covariates used in the individual models. The grey band indicates the non-significance range. The first-order step length term is consistently negative and significant, while the second-order term is positive, reflecting a characteristic step length preference. Log-transformed distance to shore fhas a strong and consistently negative effect, indicating selection for proximity to shorelines. The effect of Upland varies between individuals and is not significant on average. Palustrine Wetland shows moderate variability across individuals but is, on average, positively significant. Estuarine Wetland is positively associated with selection in nearly all individuals, though effect sizes vary.

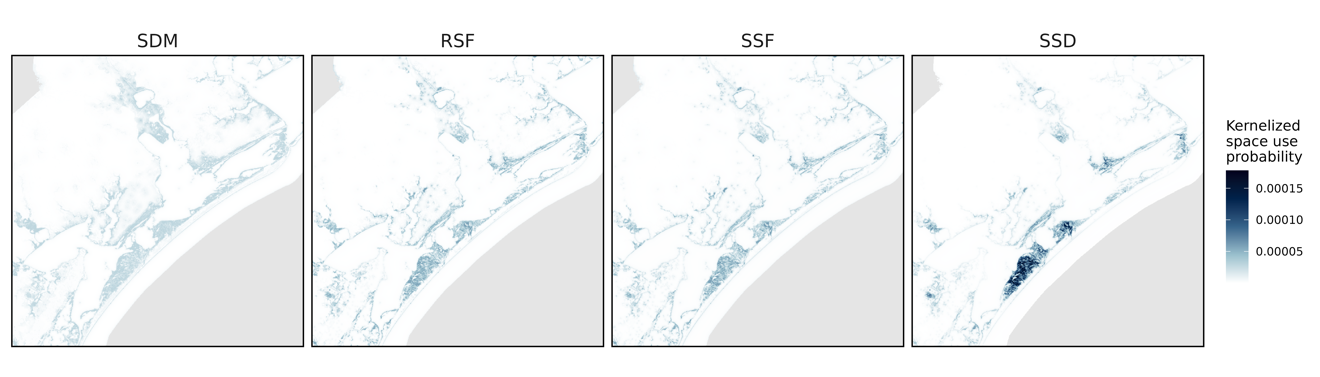

Fig. S4: Overview of spatial model predictions

Population-level predictions of crane space use probability based on SDM, RSF, SSF, and SDD models, each normalized to sum to one (150m spatial resolution).
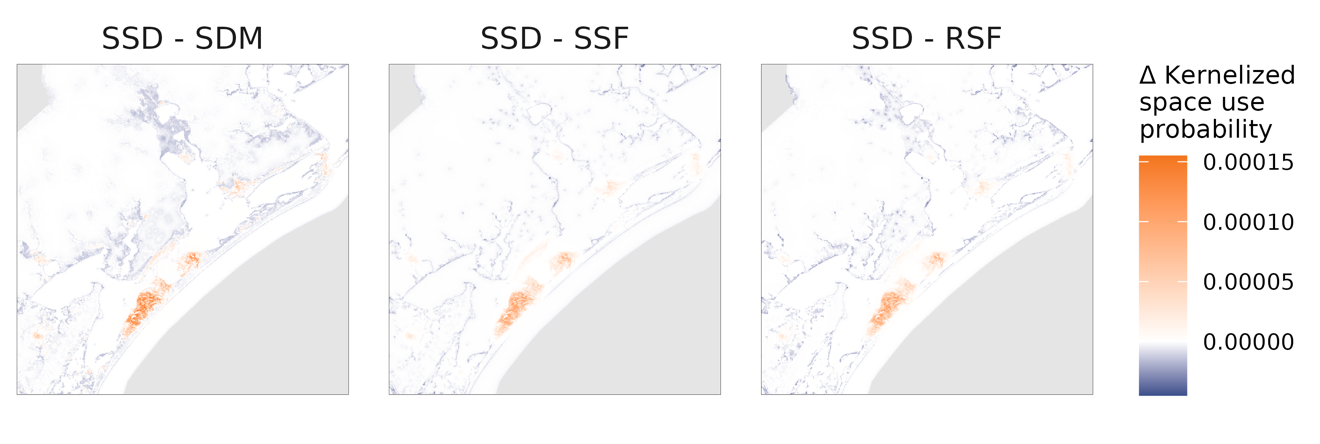

Fig. S5: Spatial differences in model predictions

Delta maps illustrate spatial variation in kernelized space use probabilities between SSD and SDM/SSF/RSF models. The panel from left to right show differences between SSD and SDM predictions (SSD - SDM), differences between SSD and SSF predictions (SSD - SSF), and differences between SSD and RSF predictions (SSD - RSF). Warm colors indicate areas where SSD predicts higher space use intensity than the respective alternative model, while cool colors highlight areas where SDM/SSF/RSF predict higher values. Compared to all other models, SSD predicts higher intensities in larger, contiguous habitat patches, whereas SDM/SSF/RSF exhibit more spatially dispersed patterns. This reflects the influence of movement constraints in the SSD model, which lead to greater spatial concentration in areas of low movement cost and longer expected retention times.

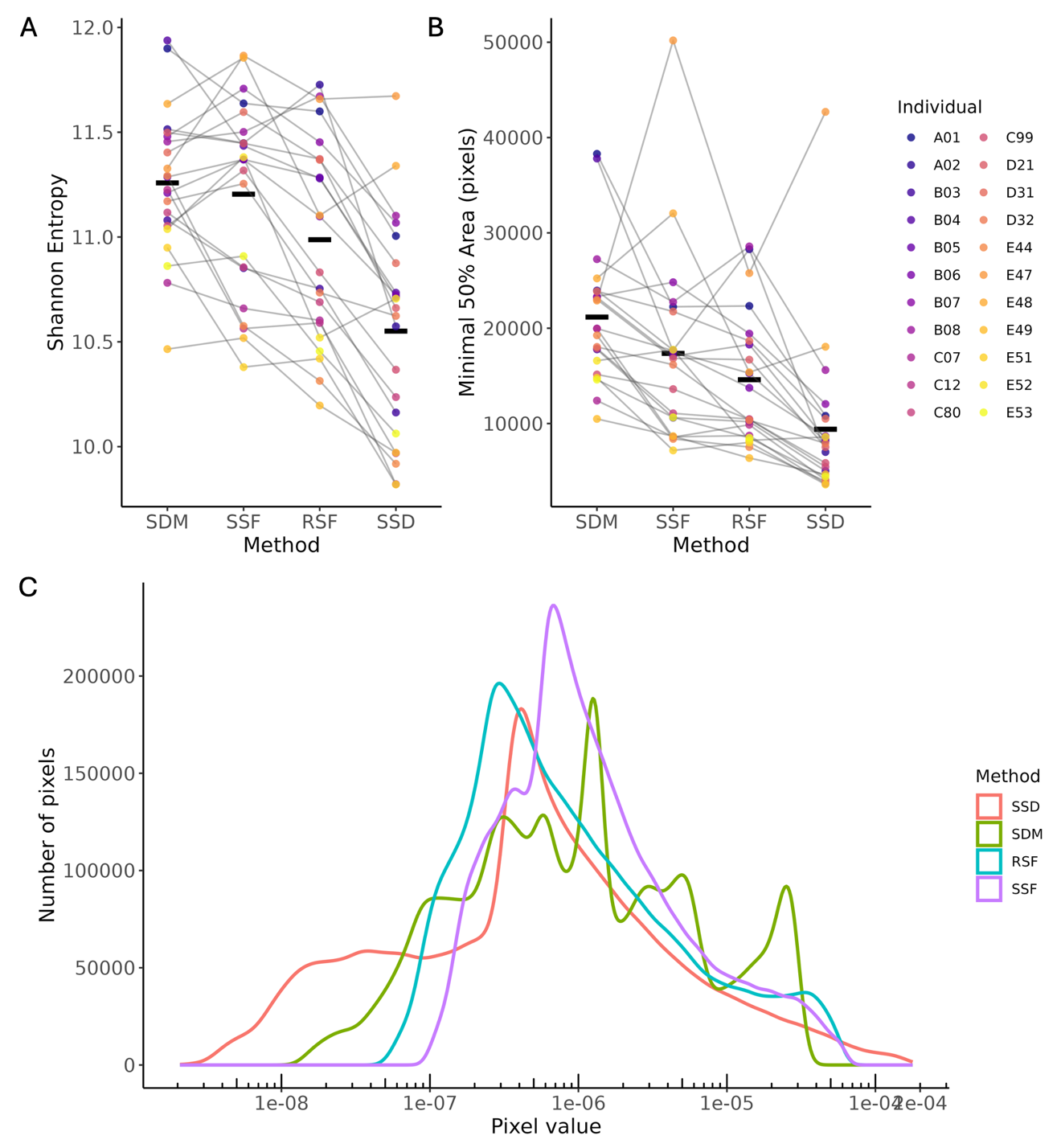

Fig. S6: Space use heterogeneity

Comparison of spatial concentration in predicted individual space use across the four modeling approaches (SDM, SSF, RSF, SSD). Two complementary metrics are shown in panels A and B to assess how evenly individuals distribute their space use across the landscape, and panel C illustrates the underlying heterogeneity in predicted use values. **(A)** Shannon entropy of individual space use distributions. Lower entropy indicates more spatially concentrated (less uniform) use, whereas higher entropy reflects more evenly distributed use. Shannon entropy declines steadily from SDM → SSF → RSF → SSD, showing that SSD predicts the most concentrated patterns. **(B)** Minimal area required to encompass 50% of an individual’s predicted space use (“min50% area”). Smaller areas indicate stronger concentration. Again, SSD yields the smallest mean area, followed by RSF, SSF, and SDM. Each colored dot represents one individual; gray lines connect the same individual across methods; black crossbars denote method means. **(C)** Kernel density curves of pixel‐value distributions from each method’s use–intensity surface. The SSD curve (red) is markedly broader, with elevated frequencies at both very low and very high pixel values, reflecting its more heterogeneous space use: SSD allocates many pixels to near‐zero use (fragmented or low‐quality areas) while concentrating use strongly in a smaller set of high‐value patches. In contrast, SDM, SSF, and RSF density curves are narrower and centered on intermediate values, indicating these methods mainly capture habitat‐quality effects without the same extremes of concentration or absence.

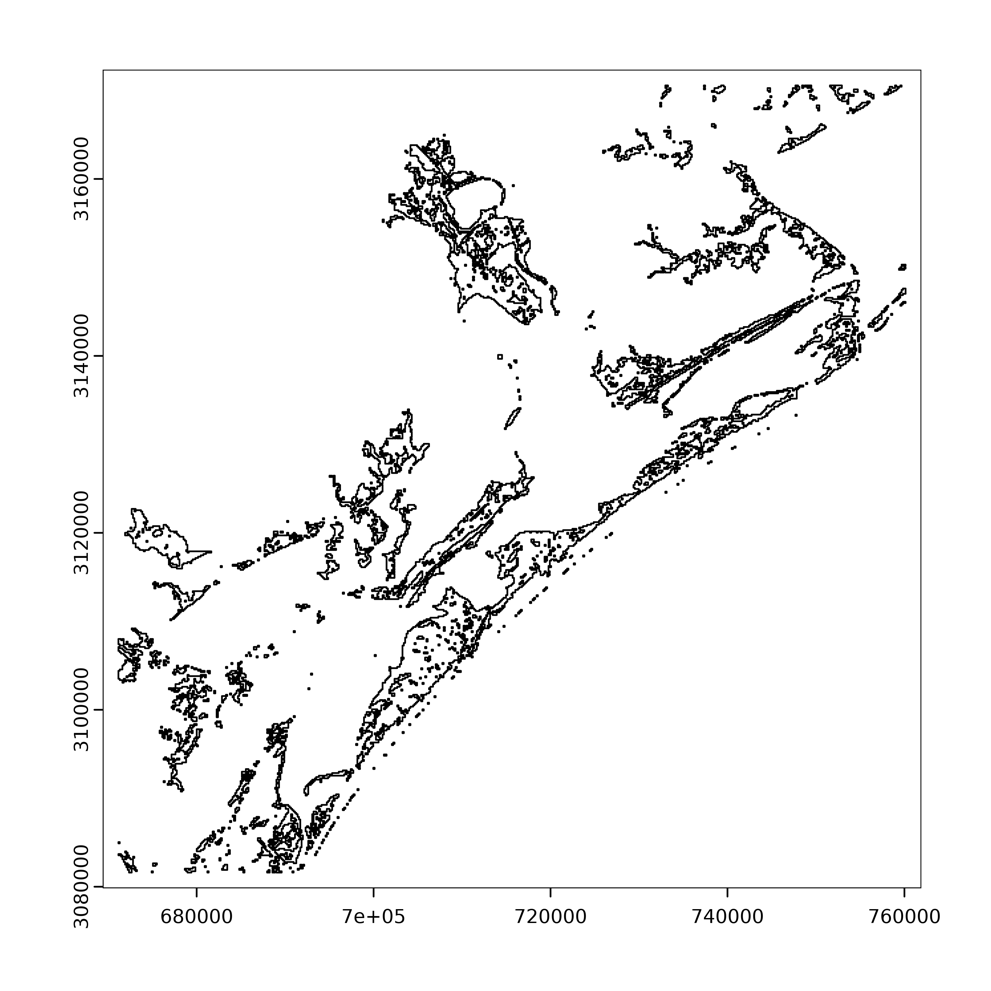

Fig. S7: Distribution of habitat patches

Polygons of the habitat patches as used in the patch-level analyses throughout the manuscript. As described in the methods section, patches are defined as contiguous areas of estuarine wetland, which is the dominant habitat type used by cranes during winter, as indicated by both model coefficients.

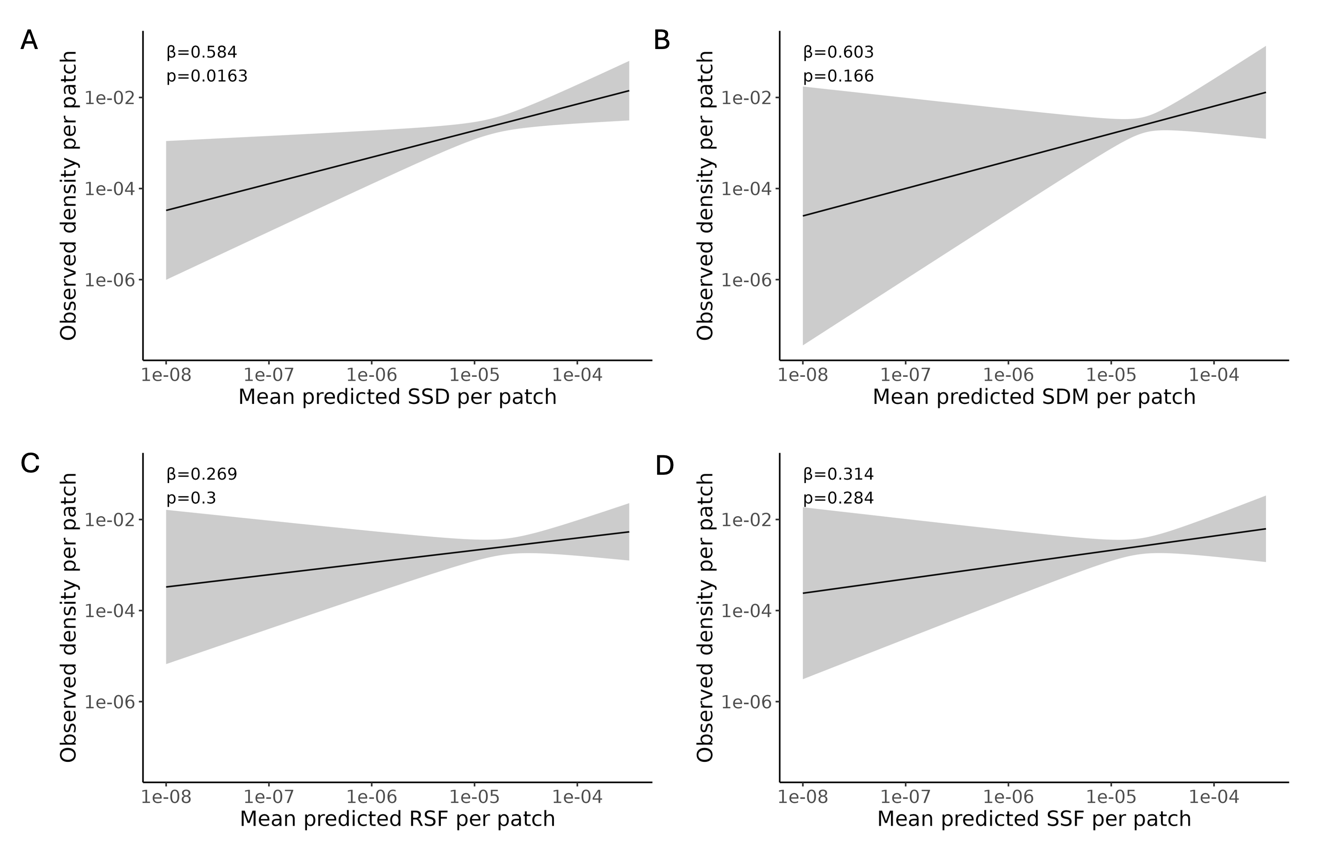

Fig. S8: Model comparisons in predicting densities

Observed patch‐level densities from independent census versus model‐predicted densities for the SSD (A), SDM (B), RSF (C), and SSF (D) models. Panels show the conditional (non‐zero) component of zero‐inflated Gamma GLMMs fit to the observed density per patch (individuals observed divided by patch size) as a function of the mean predicted value from each method. Solid black lines are the modelled relationships on log–log axes; shaded ribbons are 95 % confidence intervals. In each panel the estimated slope (β) and p‐value for the fixed effect are annotated. As highlighted in the main text, note only SSD (panel A) shows a statistically significant positive association (β = 0.584, p = 0.0163), whereas SDM, RSF and SSF (panels B–D) do not.

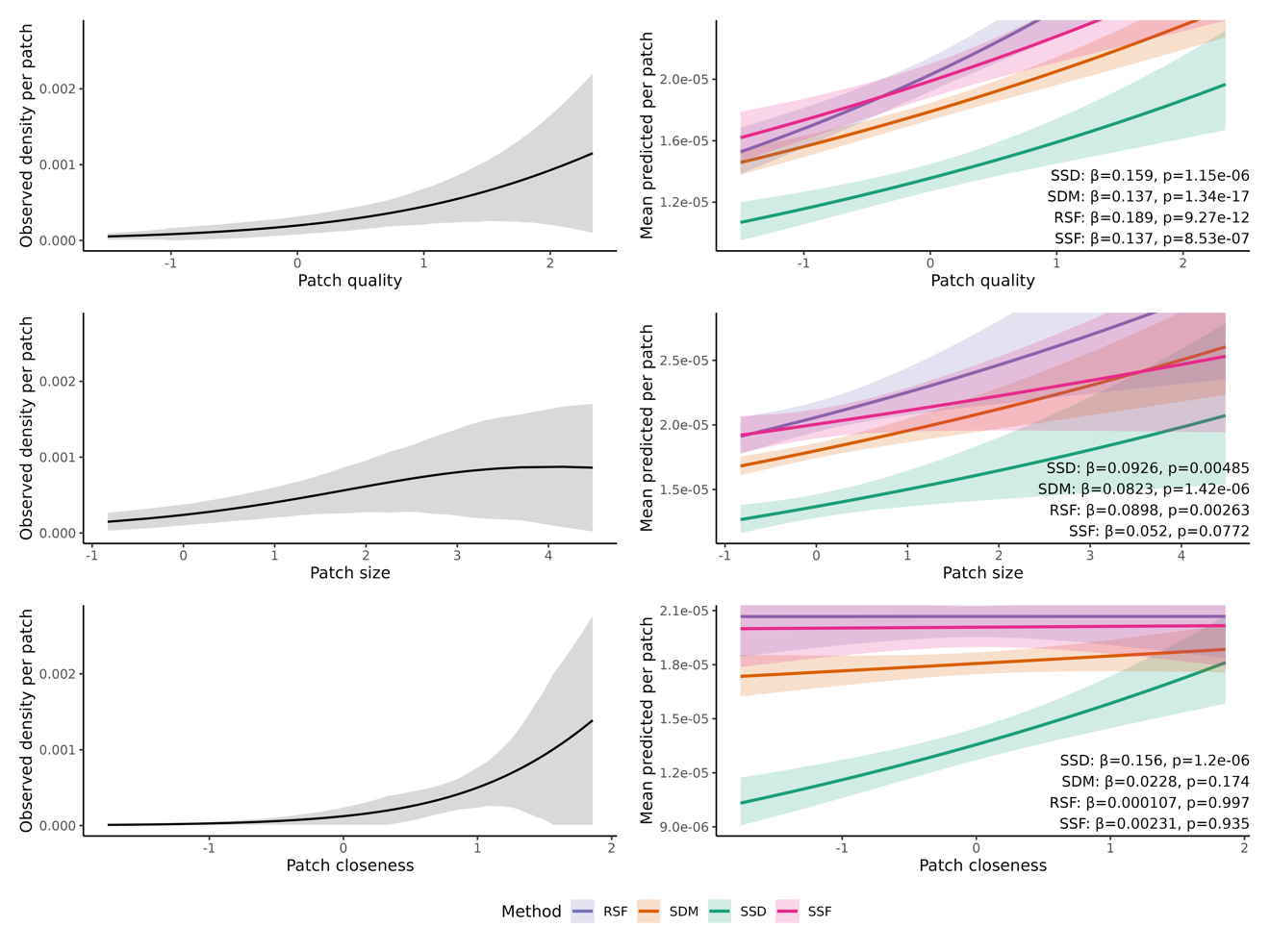

Fig. S9: Model comparisons in capturing metapopulation effects

Observed relationships between independently estimated density and patch attributes (left panels) and corresponding model-predicted values for each approach (right panels) are shown for patch quality, patch size, and patch closeness. SSD, SDM, RSF, and SSF models all capture positive effects of patch quality, consistent with their reliance on habitat preference. Patch size shows a slightly significant effect for SDM and RSF models and a stronger effect for SSDs. Only SSD models, however, additionally capture the expected significant effect of patch closeness, consistent with metapopulation theory and observed patterns. Reported β values indicate the estimated slope of respective log link Gamma GLM models, and *p*-values indicate significance. All predictors were standardized (z-score) prior to analysis.

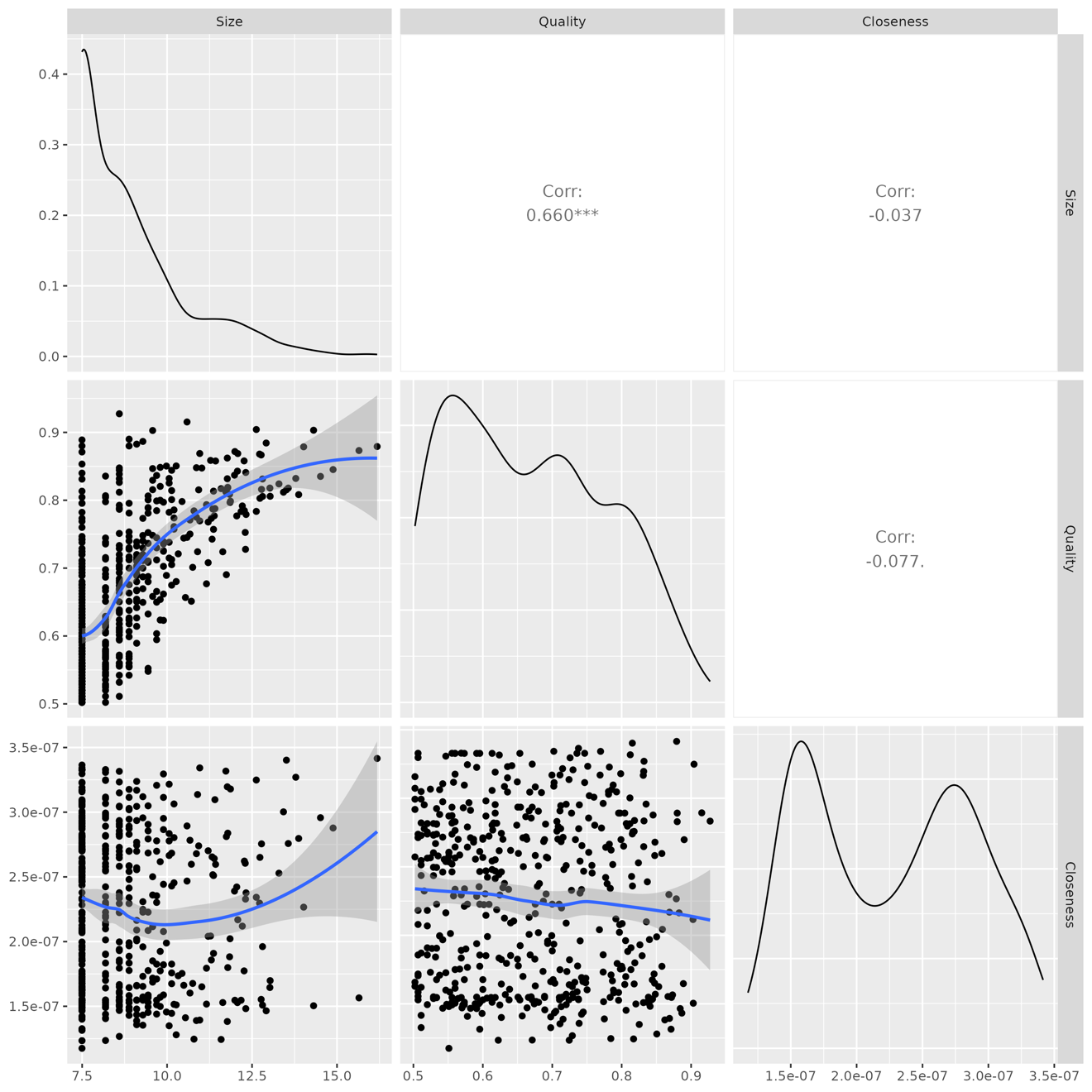

Fig. S10: Correlation analysis

Pairwise relationships among patch metrics. Scatterplot matrix showing patch size (log-transformed), habitat quality (proportion of estuarine wetland cover), and connectivity (closeness centrality). Each lower-panel scatterplot includes fitted regression lines, with histograms on the diagonal. Patch size and habitat quality are positively correlated, indicating that larger patches tend to have higher estuarine wetland cover. This correlation explains why all models appear to capture patch size effects, even though by design, only the SSD incorporates area dependence.

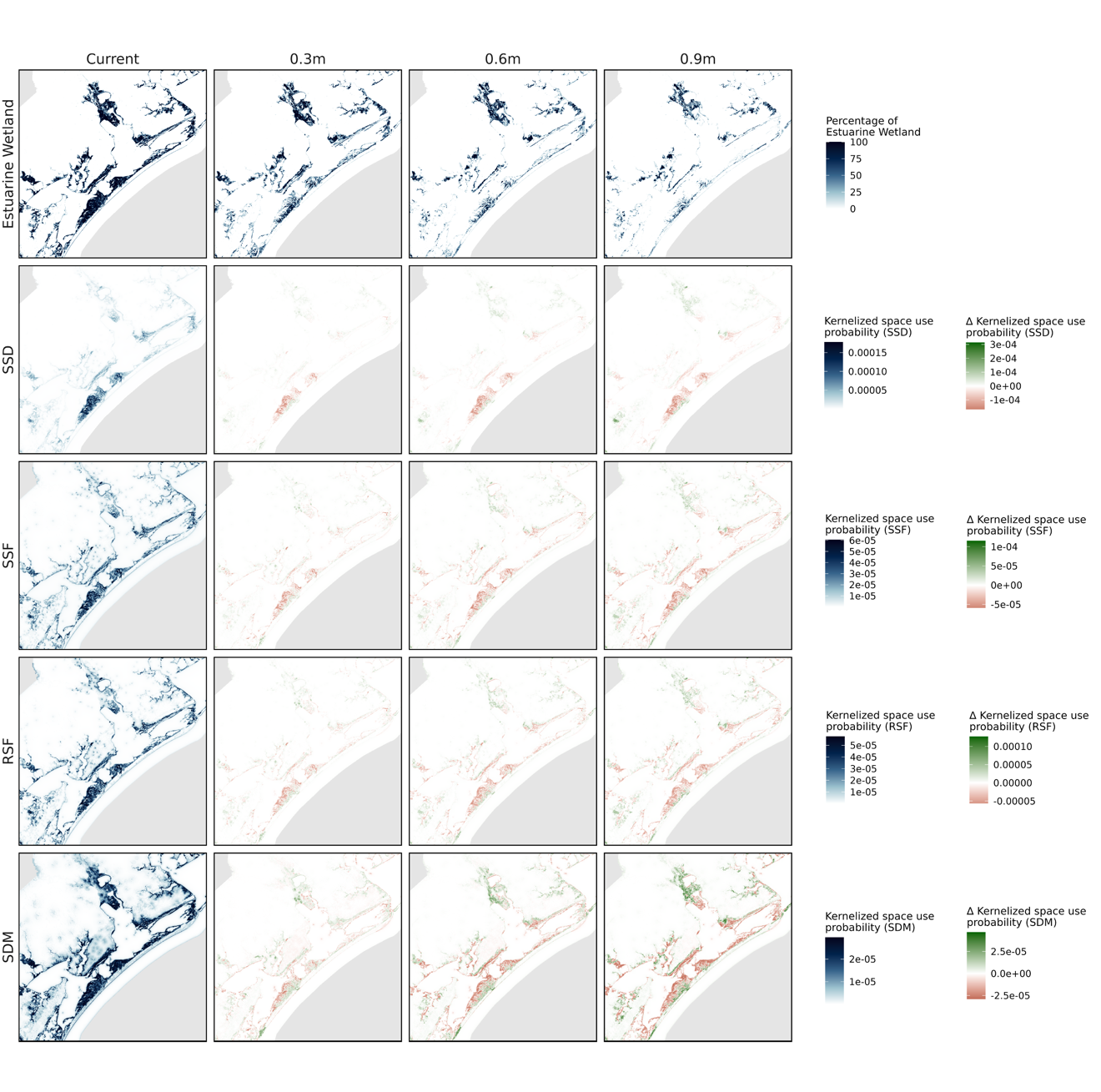

Fig. S11: Model spatial projections under future sea level rise

Spatial changes in estuarine wetlands (top row), and predicted whooping crane space use derived from SSD (second row), SSF (third row), RSF (fourth row), and SDM (bottom row) models. Columns correspond to current conditions (first column), followed by 0.3 m, 0.6 m, and 0.9 m sea-level rise scenarios. The first row displays absolute estuarine wetland cover under each scenario. The lower three rows show delta maps for each model, indicating changes in predicted space use relative to current conditions (red = decrease, green = increase). These maps illustrate how increasing sea-level rise leads to habitat loss, patch fragmentation, and model-specific shifts in predicted crane space use, with the SSD model consistently showing the most spatially constrained redistribution.

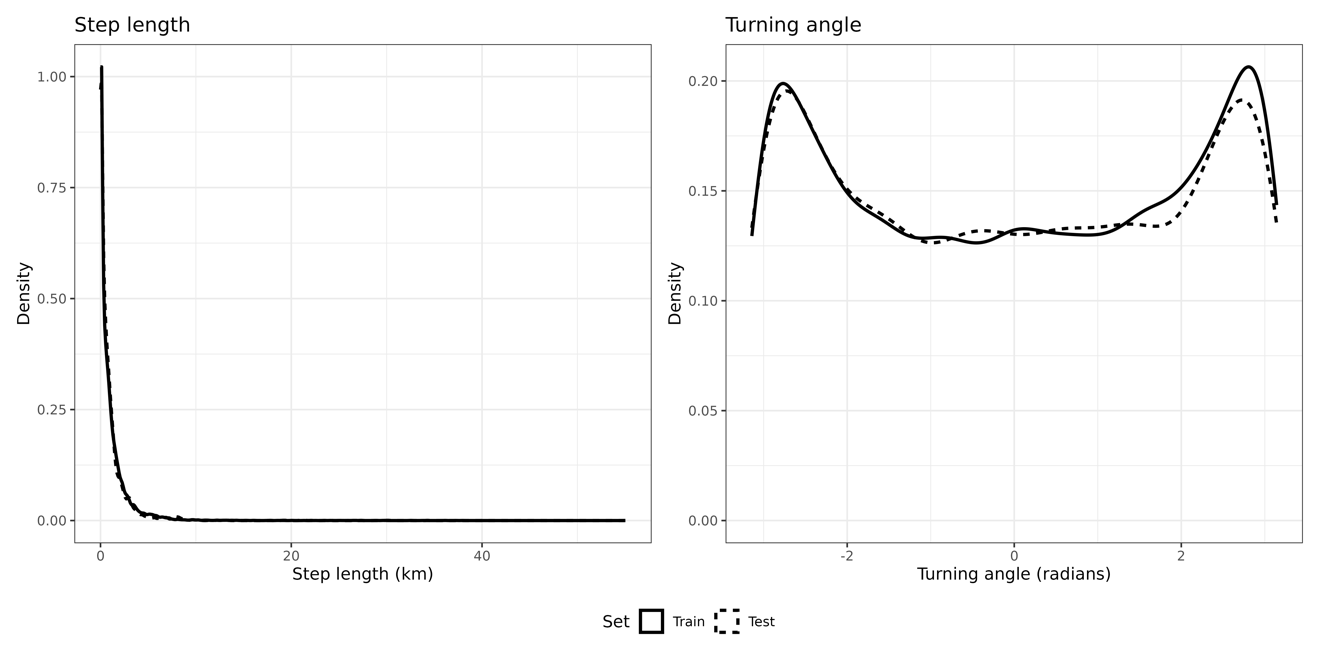

Fig. S12: Consistency of movement metrics between training and testing data

Pooled distributions of step length (left) and turning angle (right) for training (75% of data) and testing (25% of data) locations across all whooping cranes. The curves are highly similar between the two sets, indicating that movement behavior did not change systematically between training and testing subsets. This supports the assumption that splitting tracks temporally does not bias model evaluation.

Tables

Table S1: Model pseudocode

SSF models were used for both the selection-only SSF projections and the SSD surfaces. RSF models were used to generate RSF projection surfaces

| Model | Pseudocode |
| --- | --- |
| SSF | Upland + Palustrine_Wetland + Estuarine_Wetland + log(Shore_Distance + 1) + poly(sl_, 2) + strata(step_id_) |
| RSF and SDM | Upland + Palustrine_Wetland + Estuarine_Wetland + log(Shore_Distance + 1) |

*For all SSF, RSF, and SDM:*

Upland is the fraction of 10m x 10m pixels in each 150m x 150m pixel of the Upland land cover class, which is created by aggregating NOAA Marsh Migration classes "High Intensity Developed", "Medium Intensity Developed", "Low Intensity Developed", "Developed Open Space", "All Uplands";

Palustrine_Wetland is the fraction of 10m x 10m pixels in each 150m x 150m pixel of the Palustrine_Wetland land cover class, which is created by aggregating NOAA Marsh Migration classes "Palustrine Forested Wetland", "Palustrine Scrub/Shrub Wetland", "Palustrine Emergent Wetland";

Estuarine_Wetland is the fraction of 10m x 10m pixels in each 150m x 150m pixel of the Estuarine_Wetland land cover class, which is created by aggregating NOAA Marsh Migration classes "Brackish/Transition Wetland", "Estuarine Wetland", "Unconsolidated Shore";

Shore_Distance is the normalized distance to the next pixel of class "Unconsolidated Shore" in the NOAA Marsh Migration data set.

*For SSF only:*

sl_ is step length in meters;

step_id is the unique identifier of an observed step and the 100 randomly sampled points associated to it.

**SI References**

Baddeley, A., Rubak, E. & Turner, R. (2015). *Spatial Point Patterns: Methodology and Applications with R*. Chapman and Hall/CRC Press, London.

Commission for Environmental Cooperation (CEC), Statistics Canada, United States Census Bureau, & Instituto Nacional de Estadística y Geografía (INEGI). (2022). North American Atlas - Political Boundaries.

Taylor, L.N., Ketzler, L.P., Rousseau, D., Strobel, B.N., Metzger, K.L. & Butler, M.J. (2015). *Observations of whooping cranes during winter aerial surveys: 1950-2011*. Aransas National Wildlife Refuge, U.S. Fish and Wildlife Service, Austwell, Texas, USA.

UNEP-WCMC. (2025). Protected Area Profile for Wood Buffalo National Park from the World Database on Protected Areas, March 2025.

Wand, M.P. & Jones, M.C. (1994). *Kernel smoothing*. CRC press.
